# Cornichon Homolog-3 (Cnih3) deletion impairs spatial memory, reward-cue association, and fentanyl self-administration behavior

**DOI:** 10.1101/2025.11.12.688051

**Authors:** Tania Lintz, Alex Liu, Talal A. Aal, Ashley Park, Joanna J. Dearman, Arpana Agrawal, Elliot C. Nelson, Jose A. Moron

## Abstract

Opioid misuse remains rampant as new synthetic opioids reach the market. Large-scale genetic tools like the GWAS identify previously unrecognized targets and biomarkers in opioid misuse with hopes of combating the opioid epidemic. One such target is the AMPAR auxiliary protein Cornichon Homolog-3 (human analog: CNIH3, mouse analog: Cnih3), which determines AMPAR subunit composition and kinetics. Though CNIH3 was identified as a gene of interest in OUD, its role in opioid use and accompanying risk factors has not been studied. Using mice with Cnih3 deletion, we assess the role of CNIH3 in risk factors for opioid use, cognition, and opioid use itself. We find that Cnih3 deletion moderately impairs spatial memory, reward-cue association, and reversal learning. Cnih3 deletion also impairs fentanyl-cue association and blunts fentanyl intake during IVSA. We use principal component analysis to pinpoint the dimensions in which Cnih3 deletion impacts behavior in an unbiased manner. Additionally, we identify in previously published human data that single-nucleotide polymorphisms are more protective against progression to daily opioid use in women than in men, suggesting a potential sex-specific role of CNIH3. These findings highlight an important role of CNIH3 in opioid use through learning and memory processes that may differ between males and females.

## Introduction

In 2024, 2.7% of people 12 or older reported misusing opioids in the last year(*1*). Frequent use or misuse of opioids can lead to opioid use disorder (OUD), which is characterized by repeated drug-taking, drug craving during attempted abstinence, and lastly, relapse(*2*). While a decade ago, prescription opioids were the major contributors to opioid misuse, highly potent and readily available synthetic opioids, such as fentanyl, have exacerbated overdose deaths in the US(*3*). Uncovering why certain populations are predisposed to or protected against opioid misuse and OUD is critical for overdose prevention as more potent opioids continue to flood the market.

OUD is heritable, with 50-70% of variation due to segregating loci(*4*). The genome-wide association study (GWAS) is a valuable tool that can identify common variants that contribute to OUD heritability. There have been numerous large-scale GWAS of OUD (e.g.,(*5–8*)), and more recently, these GWAS have utilized control individuals with a history of some opioid exposure. Comparing OUD cases to such exposed controls ensures that identified variants specifically relate to opioid addiction rather than behavioral attributes that might contribute to using opioids initially. However, in a majority of these GWAS, exposure was minimally defined. One prior GWAS utilized a novel design in which a small group of individuals with opioid dependence who injected daily (n=1167) were contrasted with neighborhood controls, as well as a small cohort of opioid-dependent individuals who never progressed to daily injection use. Further controlling for environmental factors, controls were recruited from the vicinity of the methadone clinics where cases were recruited. This GWAS identified variants in the Cornichon-homolog 3 (human analog: CNIH3, mouse analog: Cnih3) gene (*9*). CNIH3 is an AMPAR auxiliary protein important for glutamatergic plasticity – a critical player in maladaptive drug use(*10–12*). CNIH3 regulates synaptic AMPAR expression and subunit composition necessary for long-term potentiation (LTP) by increasing surface expression of GluA1 and enhancing AMPAR kinetics(*13–15*). This form of plasticity is critical for several processes contributing to maladaptive opioid use, including learning, memory, stress-coping, nociception, and affect (*10,16–18*) with alterations found in human and animal models of major depressive disorder(*19–22*), anxiety(*23,24*), stress(*17,25*), and pain(*26,27*). Therefore, SNPs in CNIH3 may protect against OUD by disrupting plasticity that contributes to any of the above-mentioned underlying processes. That these variants arose when case status involved a severe form of OUD, and controls were individuals engaged in problematic but not severe OUD, suggests specificity of CNIH3, and given that CNIH3 determines AMPAR composition and trafficking, CNIH3 may influence *risk factors* that impact maladaptive opioid use.

Glutamate transmission is also necessary for the development, maintenance, and recall of opioid-associated memories(*10,28,29*). Alterations in CNIH3 may facilitate maladaptive opioid use by impacting the acquisition, expression, or longevity of memory, including opioid-associated memory. Increased GluA1-containing AMPARs facilitated opioid-associated contextual memory(*30–32*), and altering AMPAR subunit composition reduced opioid seeking(*33–37*). Furthermore, in previous studies, we overexpressed Cnih3 in the hippocampus and found that this enhanced the performance of female mice in the Barnes Maze spatial memory task(*38*). Our previous work also shows that female mice with a Cnih3 deletion exhibited deficits in performance on the Barnes Maze spatial memory task, as well as decreased synaptic connectivity, altered AMPAR subunit composition, and impaired LTP maintenance(*38*). Based on this evidence, CNIH3 may influence the learning and memory required for the development and progression of OUD.

Given the GWAS identification of SNPs in CNIH3 protecting against OUD, we hypothesize that CNIH3 plays an essential role in the AMPAR trafficking necessary for opioid-associated learning. To test this, we use a Cnih3 KO mouse model to assess the role of Cnih3 in risk factors to OUD (well-being, anxiety- and depression-like behavior, spatial memory, sociability, nociception), natural reward-seeking and cognitive flexibility, and last, in instrumental opioid self-administration. We found that Cnih3 deletion impairs spatial memory, the development of reward-cue associations, cognitive flexibility, and fentanyl self-administration and reinstatement. These experiments are the first to assess the role of Cnih3 in opioid use and accompanying risk factors, which seems to act through facilitating reward-associated memory formation. These results lay the foundation of CNIH3 as a promising therapeutic target.

## Results

### SNPs in CNIH3 are more protective in females against the progression to daily opioid use

In this study, we first reanalyzed data from the Comorbidity and Trauma Study, the sample for the original GWAS that implicated CNIH3 in severe opioid dependence with daily injection (*9*) to probe for sex differences. We found that, of the six SNPs in CNIH3 found to be protective against the progression to daily injection opioid use, three (rs1436171, rs1369846, and rs1436175) were more protective in women than men (Table 1). These data suggest that CNIH3-related processes (such as expression, post-translational modification, function, etc.) may differ in a way that produces sex specific effects on behavior.

**Table 1:** Evaluation of sex differences in GWAS findings from Nelson, 2016.

|  | CATS-Nelson, 2016 (n=1328) |  | Women (n=537) |  | Men (n=791) |  |
| --- | --- | --- | --- | --- | --- | --- |
| CNIH3SNP | OR (95%CI) | p-value | OR (95%CI) | p-value | OR (95%CI) | p-value |
| rs10799590 | 0.55 (0.43–0.70) | 1.28E-06 | 0.44 (0.30–0.64) | 1.60E-05 | 0.65 (0.47–0.90) | 1.03E-02 |
| rs12130499 | 0.54 (0.42–0.69) | 9.59E-07 | 0.42 (0.29–0.61) | 6.44E-06 | 0.66 (0.48–0.92) | 1.32E-02 |
| rs298733 | 0.55 (0.43–0.70) | 1.27E-06 | 0.43 (0.29–0.62) | 9.78E-06 | 0.66 (0.48–0.92) | 1.27E-02 |
| rs1436171 | 0.53 (0.42–0.68) | 5.29E-07 | 0.40 (0.27–0.58) | 2.32E-06 | 0.67 (0.48–0.93) | 1.53E-02 |
| rs1369846 | 0.51 (0.40–0.65) | 6.30E-08 | 0.38 (0.26–0.56) | 8.57E-07 | 0.65 (0.47–0.88) | 6.55E-03 |
| rs1436175 | 0.50 (0.39–0.64) | 3.15E-08 | 0.36 (0.24–0.53) | 4.49E-07 | 0.63 (0.46–0.86) | 3.92E-03 |
| p<0.05 indicating a significant interaction with greater protective effects in women |  |  |  |  |  |  |
| CATS:Comorbidity and Trauma Study, OR: OddsRatio, SNP: Single Nucleotide Polymorphism, CI: ConfidenceInterval |  |  |  |  |  |  |

### Cnih3 deletion does not impact well-being, motor coordination, or baseline nociception

To fully assess the potential role of CNIH3 in opioid use and associated risk factors, we employ a preclinical mouse model using a global Cnih3 deletion(*38*). First, we conducted a battery of behavioral tests to assess the role of Cnih3 in overall well-being, motor coordination, and nociception, which may impact the ability to perform operant tasks. First, we measured well-being using the nest-building assay, motor coordination using the accelerating Rotarod, and thermal nociception using the hot plate assay (Fig 1A). There were no differences between sexes or genotype in untorn nestlet (Fig 1B), latency to fall from the Rotarod on the first (Fig S1) or last trial (Fig 1C), or paw withdrawal latency on the hot plate test (Fig. 1D) indicating that Cnih3 deletion does not have baseline effects on well-being, motor coordination, or thermal nociception. Together, these findings demonstrate that Cnih3 does not contribute to innate nesting behavior or sensorimotor function.

**Figure 1:**
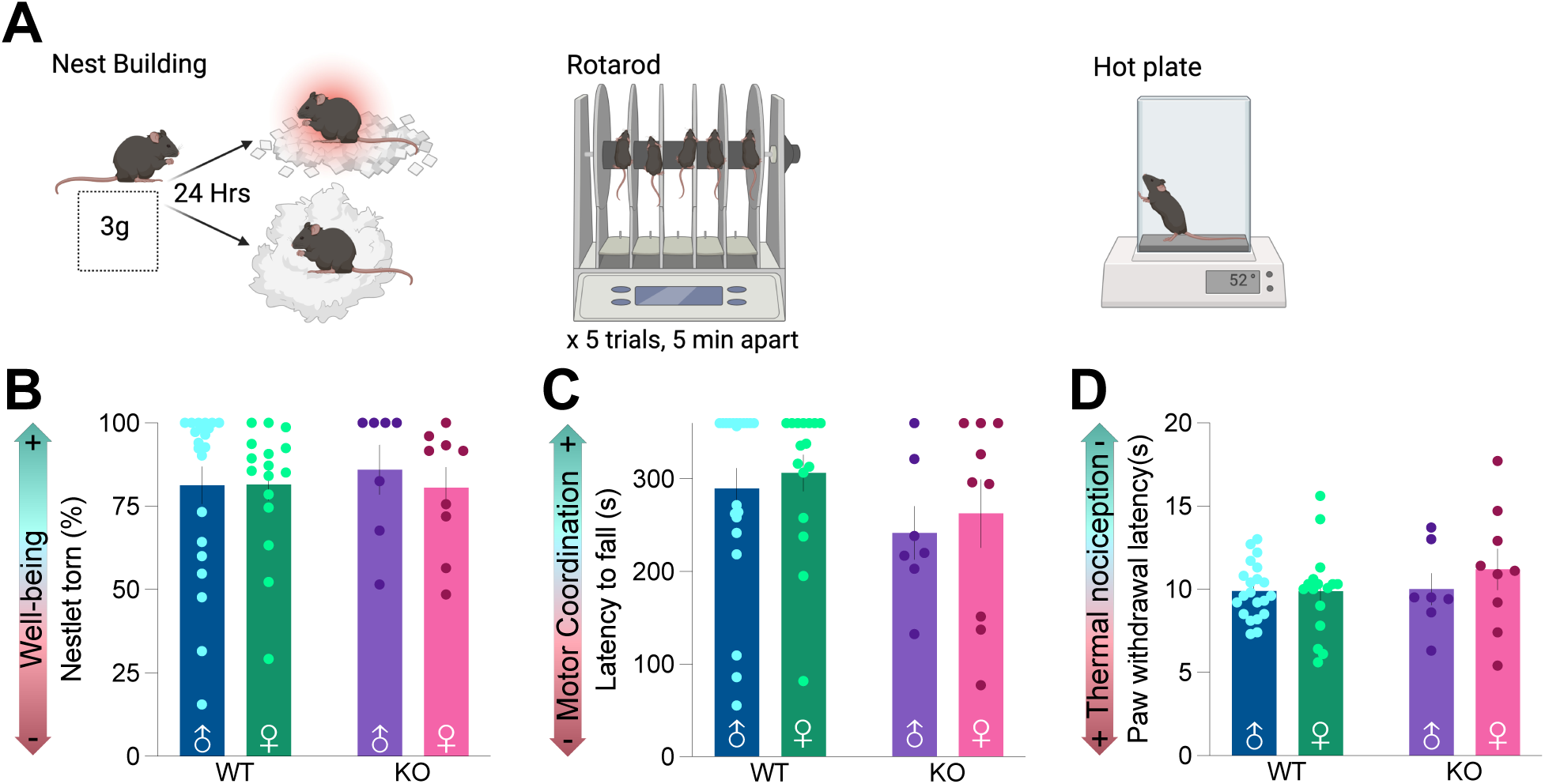
Cnih3 deletion does not impact well-being, motor coordination, or baseline nociception. A. Schematic of nest building, rotarod, and hot plate experiments. B. Well-being as measured by percentage of nestlet torn during the 24-hr testing period in male (♂) and female (♀) wildtype (WT) and Cnih3 knockout (KO) mice. C. Motor coordination as measured by latency to fall (s) on the last trial of the accelerating rotarod. D. Thermal nociception as measured by latency (s) to paw withdrawal on the hot plate set to 54°C. Data are shown as mean ± SEM. Data analyzed with two-way ANOVAs (sex x genotype). Full statistical reporting is available in Table S1. Additional data related to this dataset are available in Figure S1.

### Cnih3 deletion impairs spatial memory but does not affect socialization or social memory

Next, given the importance of associative memory in OUD, and prior work showing spatial memory deficits in Cnih3 knockout mice(*38*), we assessed whether Cnih3 KO led to memory impairments. First, we assessed spatial memory using the Novel Object Recognition Test (NORT; Fig 2A) in which mice were exposed to an open field under different conditions for 10 min/day. The first day (D1), mice were placed in an empty open field to habituate them to the apparatus and examine exploratory behavior (Fig. 2B). On day two (D2), mice returned to the field and were allowed to explore two identical objects. On day 3 (D3), one of the objects was replaced with a novel object, and the latency and total time investigating the novel object were measured as indices of spatial memory (Fig. 2C-D). Interestingly, independent of sex, Cnih3 deletion decreased total investigation time (Fig 2B), increased latency to investigate the novel object (Fig 2C), and reduced time investigating the novel object (Fig 3D), suggesting that Cnih3 may be important for spatial memory, consistent with our prior work(*38*). Next, we assessed sociability and social memory using the social interaction test (SIT; Fig 2E), in which mice were exposed to a three-chambered apparatus under different conditions for 10 min/day. On the first day (D1), mice were placed in the empty apparatus (including an empty cup in the left- and right-most chambers) to habituate them to the apparatus and assess exploratory behavior (Fig 2F). On day two (D2), mice returned to the apparatus and were allowed to explore a cup containing an unknown mouse, M1, or an empty cup to assess socialization (Fig 2G). On day 3 (D3), the previously empty cup contained a novel mouse, M2, while the familiar mouse (M1) remained on the opposite side. Time investigating M2 was measured as an indication of social memory (Fig 2H). Cnih3 deletion increased total investigation time in both sexes on D1 (Fig 2F), but there were no group differences in time investigating M1 on D2 (Fig 2G) or time investigating M2 on D3 (Fig 2H), indicating that Cnih3 does not impact social behavior or memory. Together, these findings show that Cnih3 deletion impairs spatial memory in the context of external cues, but not social factors.

**Figure 2:**
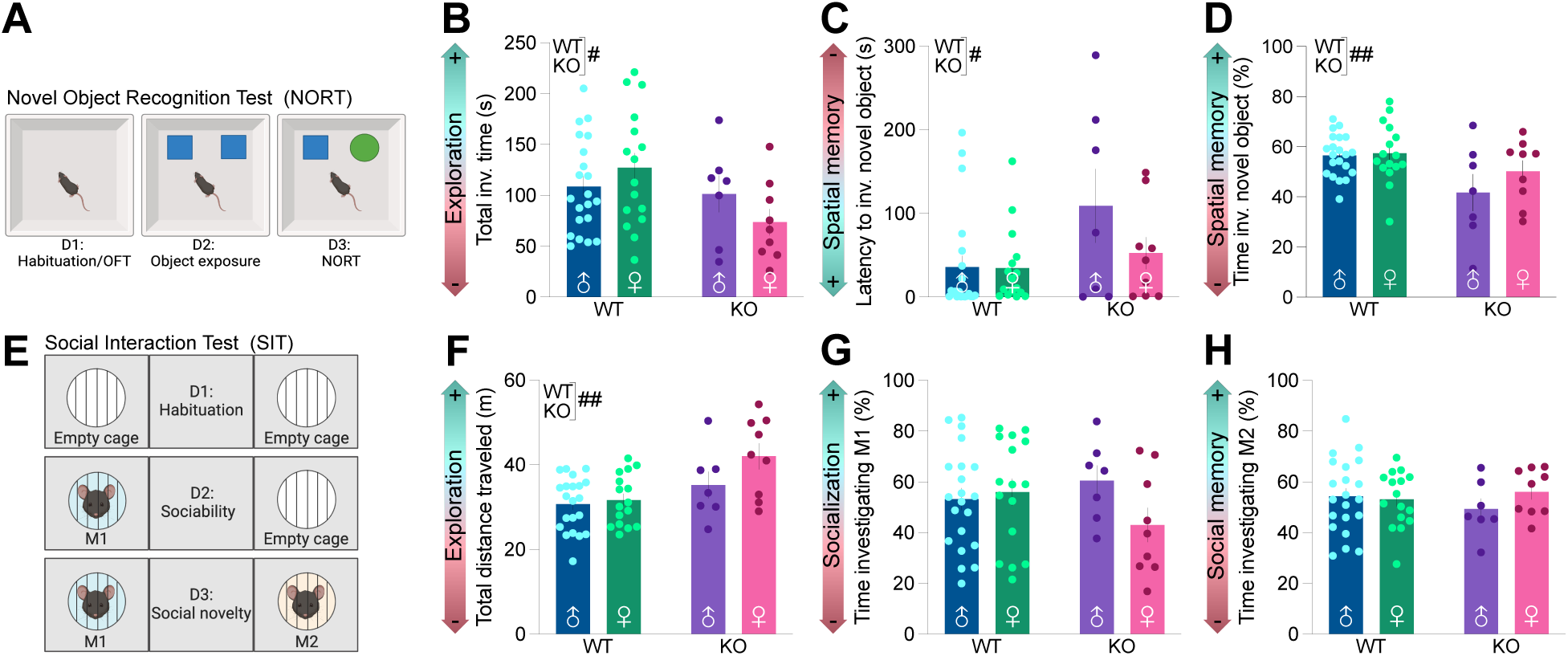
Cnih3 deletion impairs spatial memory but does not affect socialization or social memory. A. Schematic of the Novel Object Recognition Test (NORT). D1: Habituation/OFT, D2: Object exposure, D3: NORT B. Exploratory behavior as measured by total investigation time (s) during NORT D1. C. Spatial memory as measured by latency (s) to the first investigation of the novel object on NORT D3. D. Spatial memory as measured by time investigating (s) the novel object on NORT D3. E. Schematic of the Social Interaction Test (SIT). D1: Habituation, D2: Sociability, D3: Social novelty, M1: known mouse, M2: novel mouse. F. Exploratory behavior as measured by total distance traveled (m) during SIT D1. G. Socialization as measured by time investigating (%) M1 on SIT D2. H. Social memory as measured by time investigating (%) M2 on SIT D3. Data are shown as mean ± SEM. Data analyzed with two-way ANOVAs (sex x genotype). Symbols denote **^#^**main effect of genotype (**^#^**p < 0.05, **^##^**p < 0.01). Full statistical reporting is available in Table S1.

**Figure 3:**
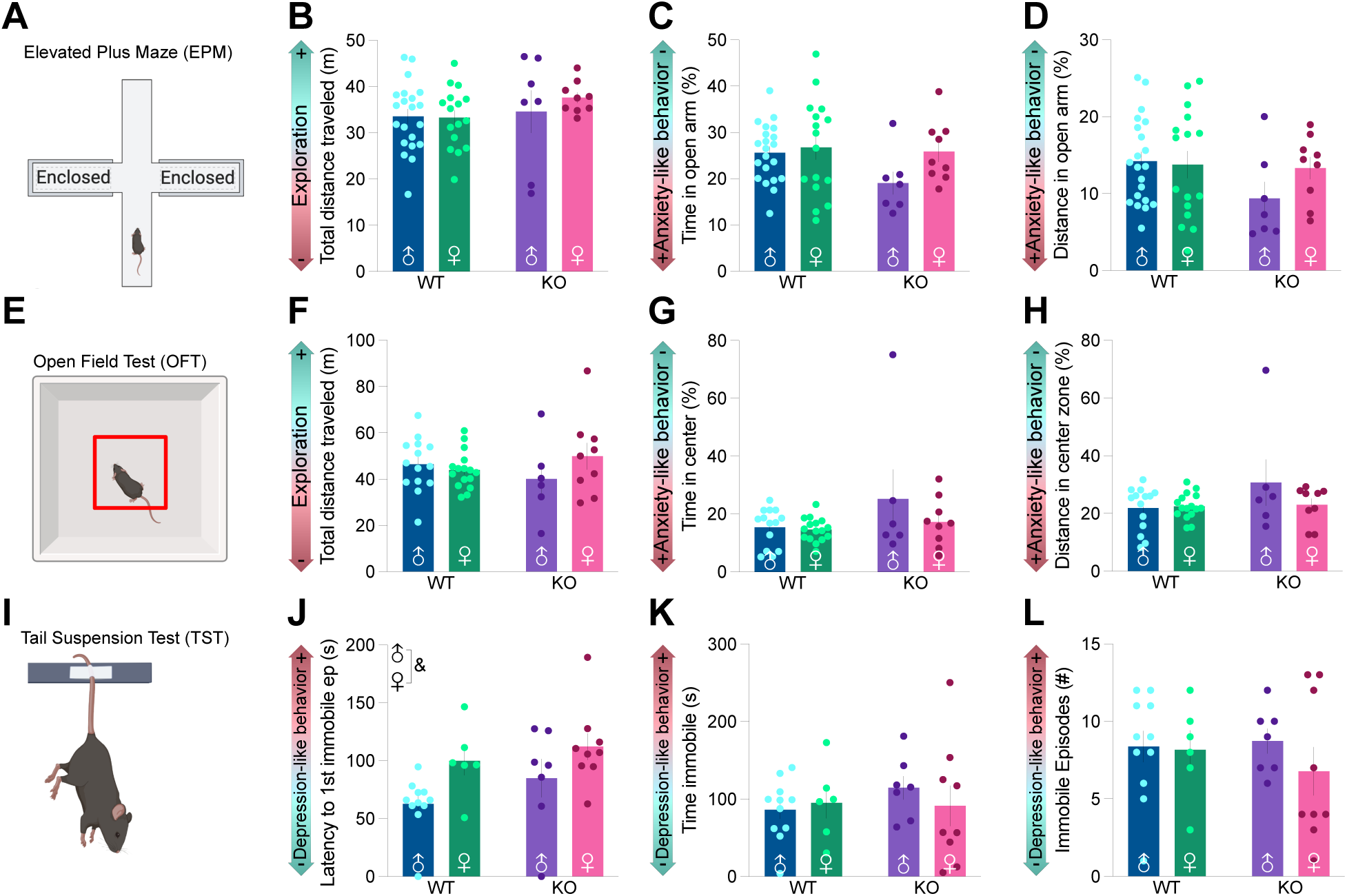
Cnih3 deletion does not significantly impact affective behaviors. A. Schematic of the Elevated Plus Maze (EPM) experiment. B. Exploratory behavior during EPM as measured by total distance traveled (m) during EPM. C. and D. Anxiety-like behavior as measured by C. time spent in the open arm (%) and D. distance traveled in open arm (%) during EPM. D. Schematic of the Open Field Test (OFT) experiment. E. Exploration as measured by total distance traveled (m) during OFT. F. and H. Anxiety-like behavior as measured by G. time in the center zone (%) and H. distance traveled in the center zone (%) of OFT. I. Schematic of the Tail Suspension Test (TST). J., K. and L. Depression-like behavior as measured by J. latency to the first immobile episode, K. total time immobile (s) and L. number of immobile episodes during the TST. Data are shown as mean ± SEM. Data analyzed with two-way ANOVAs (sex x genotype). Symbols denote ***^&^***main effects of sex in J (***^&^***p < 0.05). Full statistical reporting is available in Table S1. Additional data related to this dataset are available in Figure S2.

### Cnih3 deletion does not significantly impact affective behaviors

We next assessed how Cnih3 deletion impacted affective behaviors, which are common risk factors for OUD (*39–41*). We assessed anxiety-like behavior using the Elevated Plus Maze (EPM; Fig 3A) in which mice were placed in the center of the 4-armed apparatus (2 open, anxiogenic arms; 2 enclosed, anxiolytic arms) and allowed to explore for 10 minutes. We observed no differences between groups in exploratory behavior measured by total distance traveled in the maze (Fig 3B) or anxiety-like behavior measured by time spent (Fig 3C) and distance traveled (Fig 3D) in the open arms. We also probed anxiety-like behavior and exploration using the Open Field Test (OFT; Fig 3E) in which mice freely explored a square apparatus for 10 minutes. Similar to the EPM, we observed no differences between groups in total distance traveled in the apparatus (Fig 3F) or anxiety-like behavior indicated by time spent (Fig 3G) and distance traveled (Fig 3H) in the anxiogenic, center zone. Last, we measured learned helpless behavior, a model of depression-like behavior, using the Tail Suspension Test (TST; 3I) in which mice were suspended by their tail for 6 minutes and latency to the first immobile episode (Fig 3J), time spent immobile (Fig 3K), and the number of immobile episodes (Fig 3L) were measured. We observed sex differences in latency to the first immobile episode, with females taking longer to reach immobility (Fig 3J), but no genotype differences. We saw no sex or genotype effects on time spent immobile (Fig 3K) or number of immobile episodes (Fig 3L). Together, these results indicate that Cnih3 deletion does not impact anxiety-like or depression-like behavior, suggesting that Cnih3 does not modulate affective behaviors.

### Cnih3 deletion has a modest effect on spatial memory and affect

To obtain an unbiased and comprehensive view of Cnih3 involvement in baseline behaviors and risk factors for OUD, we ran a principal component analysis (PCA) on the data presented in Figures 1-3. We used the PCA to identify strong patterns within this large dataset to observe which principal components (PCs) contributed the most variance in the dataset based on genotype, with PC1 and PC2 being the largest contributors. To visually assess similarity between groups, we plotted PC1 scores on the x-axis and PC2 scores on the y-axis for each mouse (each dot corresponds to one mouse) in Fig 4A. Next, we assessed group differences in the largest contributors of dataset variance by comparing PC1 (Fig 4B) and PC2 (Fig 4C) loadings between sex and genotype. Variables included in the analysis were clustered by their intended measure: well-being (Nest building, rotarod, HP), spatial/social memory (NORT, SIT), and affect (EPM, OFT, TST) (see complete list in Fig S3A), and obtained the absolute value (abs) of the loading of each measure for PC1 (Fig 4D) and PC2 (Fig 4E). Loadings of 0.3 (dotted line) and above were considered to have a significant contribution, with values approaching 1 indicating a greater contribution. Finally, the summary for each PC was calculated by averaging the PC loadings (abs) for each variable in the corresponding category. Interestingly, the PC scores revealed no clear group clusters, indicating only moderate genotype differences in this dataset (Fig 4A), with PC1 and PC2 accounting for a cumulative 29.14% of the variance (Fig S3B). For PC1, WT males and Cnih3 KO females had positive loadings, while WT females showed slightly negative, and KO males showed very negative loadings (Fig 4B). The largest loadings, and therefore largest contributors, to genotype-based variance in PC1 correspond to spatial/social memory and affect (Fig 4D), suggesting that Cnih3 plays the largest role in these traits. For PC2, WT mice had positive loadings while KO mice had negative loadings (Fig 4C). The largest loadings, and therefore largest contributors, to genotype-based variance in PC2 correspond to affect (Fig 4E), suggesting that Cnih3 plays the largest role in these traits. Taken together, these data show that there are moderate differences between genotypes in the risk factors for OUD, and that these differences are rooted in spatial memory and affect. Generally, according to PC1, WT male and KO female mice present with the following slight changes: long latencies to fall from the Rotarod, low time investigating the known object, high latency to investigate the novel object, high distance traveled on D1 of the SIT, high times spent in the open arm of the EPM, and low distance in the center zone and high distance in the center zone of the OFT, while KO females present with the opposite. According to PC2, WT males present with low amounts of untorn nestlet, high paw withdrawal on the HP, and high amounts of time spent with the empty cup on SIT D2 and both M1 and M2 on SIT D3. They also show low distance traveled on the open arm of the EPM, and low time in the center zone of the OFT, and low numbers of immobile episodes on the TST, while KO mice exhibit the opposite.

**Figure 4:**
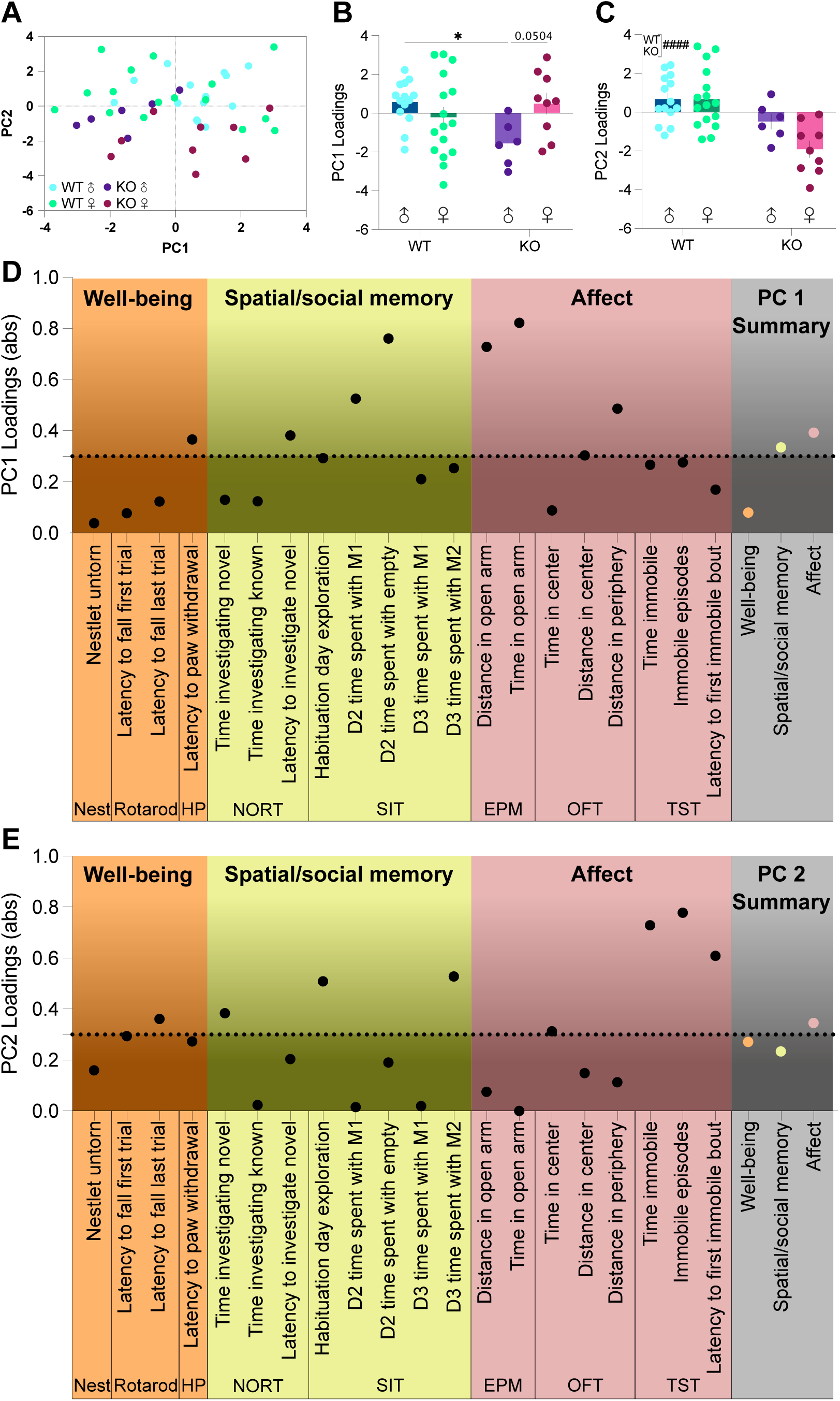
Cnih3 deletion has a modest effect on spatial memory and affect. A. Loadings plot of PC1 and PC2 for each mouse that completed the battery of behaviors described in Figures 2-4 (n=45). B. PC1 loadings for each mouse. C. PC2 loadings for each mouse. D. PC1 loadings of the variance each described behavior contributes to the dataset. The PC1 summary is the average of the PC loading of all components within a defined type of measure. Loadings above 0.3 are considered significant contributors. E. PC2 loadings of the variance each described behavior contributes to the dataset. The PC2 summary is the average of the PC loading of all components within a defined type of measure. Loadings above 0.3 are considered significant contributors. Data are reported as mean ± SEM in B-C. Average loading of each measure for the dataset shown in D-E. Two-way ANOVA (sex x genotype) and Sidak’s multiple comparisons in B and C. Symbols denote *post-hoc group differences in B (*p < 0.05) and a ***^#^***main effect of genotype in C (***^####^***p < 0.0001). Full statistical reporting is available in Table S1. Additional data related to this dataset are available in Figure S3.

### Cnih3 deletion impairs reward-cue association in females and spatial reversal learning in males

We demonstrated that Cnih3 deletion impairs spatial memory (Fig 2). To further probe the effects of Cnih3 deletion on memory and cognitive flexibility, we ran an operant sucrose self-administration and spatial reversal learning task (Fig 5A). Mice were trained to poke a light-cued nose poke port to receive a chocolate-flavored sucrose pellet during the cue-association phase. Time to reach acquisition (Fig 5B; 3 consecutive sessions where at least 70% of pokes were on the reward-producing port and the session maximum of 30 pellets were obtained), as well as rewards obtained (Fig 5E) and nose poke discrimination on the first three days (Fig 5H) and the last day (Fig 5K) of cue-association were assessed to probe learning. Then, mice underwent the cue-disassociation phase, where they poked the uncued port to receive sucrose pellets (Fig 5C-L), and the cue-reassociation phase, where they poked the cued port to receive rewards, (Fig 5D-J) to probe cognitive flexibility. During cue-association, Cnih3 KO females took longer to reach acquisition criteria compared to WT females or Cnih3 KO males (Fig 5B), which likely contributed to session interactions in rewards earned (Fig 5E) and apparent discrimination (Fig 5H) during the first 3 days of this phase, and sex differences in discrimination on the last day (Fig 5K). During the cue-disassociation phase, we saw sex but not genotype differences, with females taking longer to reach acquisition criteria (Fig 5C) and a trend for a sex by genotype interaction for rewards (Fig 5F), but no differences in discrimination, on the first three days. However, on the last day of cue-disassociation, KO mice discriminated better than their WT counterparts (Fig 5L). Last, during cue re-association, Cnih3 deletion led to increased sessions to acquisition compared to WT males and a trend for increased sessions to acquire compared to KO females (Fig 5D). There were, however, no differences in rewards earned (Fig 5G), with a sex by genotype interaction for discrimination (Fig 5J) on the first three days of cue re-association. On the last day of cue-reassociation, Cnih3 deletion produced increased discrimination in males, with no difference in females (Fig 5M). Together, these data show that Cnih3 deletion impairs initial reward-cue associations in females and cognitive flexibility as measured in our reversal learning task in males.

**Figure 5:**
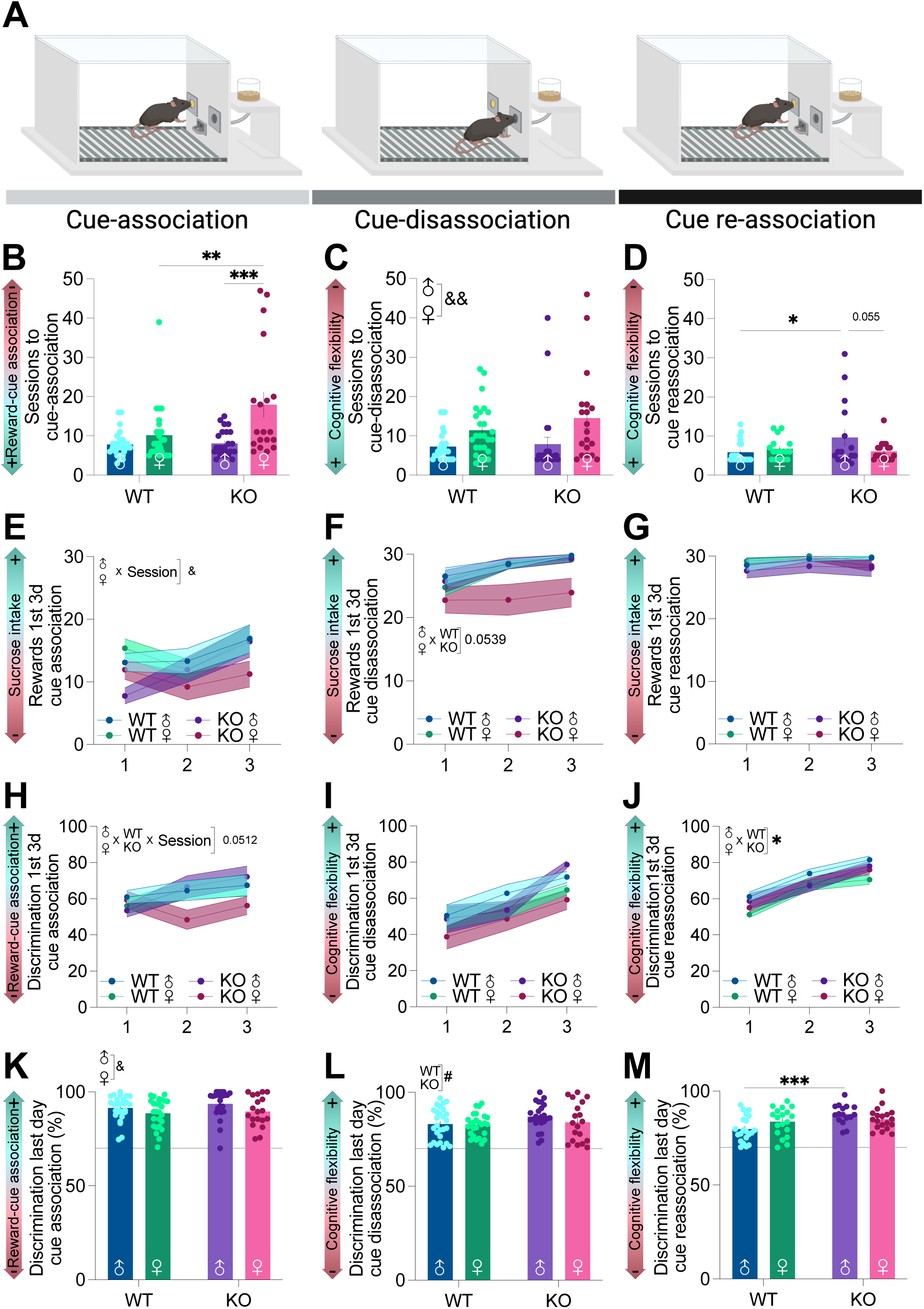
Cnih3 deletion impairs reward-cue association in females and spatial reversal learning in males. A. Experimental schematic of the sucrose self-administration/spatial reversal learning assay. B., C., and D. B. Reward-cue association or C. and D. cognitive flexibility as measured by the number of sessions to reach the acquisition criteria in B. cue-association, C. cue-disassociation, and D. cue-reassociation. E., F., and G. E. Reward-cue association or F. and G. cognitive flexibility as measured by the number of rewards earned on the first three days of E. cue-association, F. cue-disassociation, and G. cue-reassociation. H., I., and J. H. Reward-cue association or I. and J. cognitive flexibility as measured by discrimination of the reward-producing nose poke (% of pokes) on the first three days of H. cue-association, I. cue-disassociation, and J. cue-reassociation. K., L., and M. L. Reward-cue association or L. and M. cognitive flexibility as measured by discrimination of the reward-producing nose poke (% of pokes) on the last day of K. cue-association, L. cue-disassociation, and M. cue-reassociation. Data are reported as mean ± SEM. Two-way ANOVA (sex x genotype) in B-D, K-M. Symbols denote main effects of ***^#^***genotype (***^#^***p<0.05) or ***^&^***sex (***^&^*** p<0.05, ***^&&^***p<0.01) and *Sidak’s multiple comparisons (*p < 0.05, ** p < 0.01, *** p < 0.001). Three-way ANOVA (sex x genotype x session) in E-J. Symbols denote effects of ***^&^***sex x session (***^&^*** p<0.05) and *sex x genotype (*p < 0.05). Full statistical reporting is available in Table S1, and additional data are available in Figure S6.

### Cnih3 deletion produces differences in reward-cue association and cognitive flexibility

To assess the impact of Cnih3 deletion on natural reward-seeking and spatial reversal learning, we ran a PCA (as in Fig. 4) on the behaviors presented in Figure 5 and observed only moderate genotype differences in this dataset (Fig 6A), with PC1 and PC2 accounting for 40.94% of the variance (Fig S4B). Variables included in the analysis were clustered by their intended measure: reward-cue association, cognitive flexibility, and cognitive inflexibility (see complete list and direction of variable loading in Fig S4A). Cnih3 deletion produced a negative PC1 loading, while WT mice produced a positive PC1 loading, suggesting that the variables comprising PC1 are negatively correlated with WT mice and positively correlated with KO mice (Fig 6B). In contrast, female PC2 loadings were more negative than those of males, independent of genotype (Fig 6C), suggesting that the variables in PC2 correlate positively in males and negatively in females. Variable clustering within each PC revealed that the largest contributors to genotype-based variance in PC1 corresponded to reward-cue association and cognitive flexibility (Fig 6D), while variance in PC2 corresponded to cognitive flexibility (Fig 6E), suggesting that Cnih3 plays the largest role in these traits in each PC. Taken together, these data show that there are moderate differences between genotypes in natural reward-seeking and cognition, rooted in reward-cue association and cognitive flexibility. Generally, knockout mice have positive PC1 loadings, meaning they tend to exhibit the following moderate differences: long acquisition times as well as few rewards and few inactive pokes on the first day of cue association, long acquisition times, high rewards, and low inactive pokes during cue disassociation, and low acquisition times, and high rewards during cue reassociation, while WT mice with negative PC1 loadings show the opposite. Male mice tend to have positive PC2 loadings, meaning they tend to exhibit slightly shorter acquisition times on the first day of cue association, short acquisition times and low inactive pokes during cue disassociation, and high acquisition times, high rewards, and low inactive pokes during cue reassociation, while female mice with negative PC1 loadings show the opposite.

**Figure 6:**
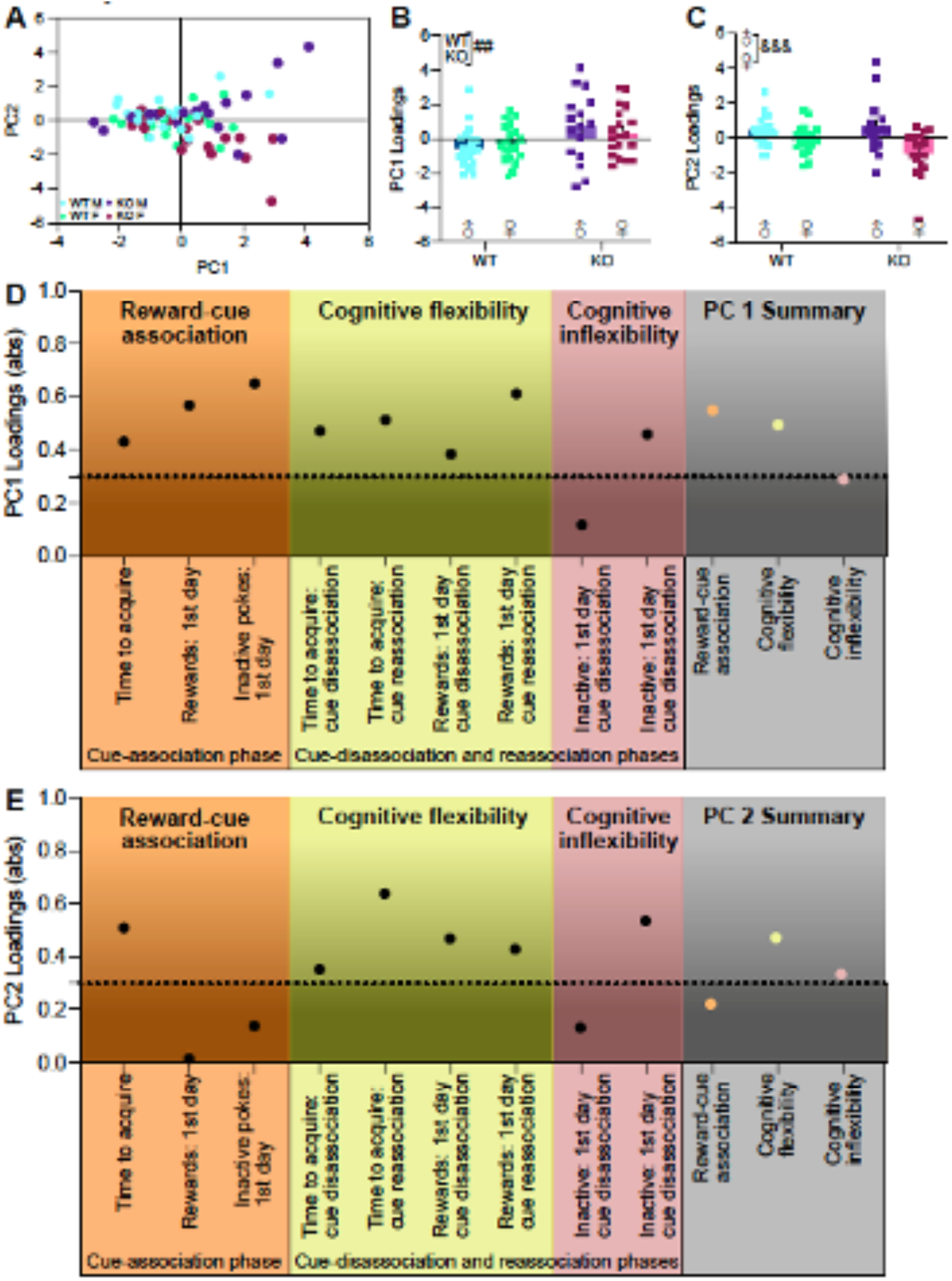
Cnih3 deletion produces differences in reward-cue association and cognitive flexibility. A. Loadings plot of PC1 and PC2 for each mouse that has completed the experiments described in Figure 6 (n=75). B. PC1 loadings for each mouse. C. PC2 loadings for each mouse. D. PC1 loadings of the variance each described behavior contributes to the dataset. The PC1 summary is the average of the PC loading of all components within a defined type of measure. Loadings above 0.3 are considered significant contributors. E. PC2 loadings of the variance each described behavior contributes to the dataset. The PC2 summary is the average of the PC loading of all components within a defined type of measure. Loadings above 0.3 are considered significant contributors. Data are reported as mean ± SEM in B-C. Average loading of each measure for the dataset shown in D-E. Two-way ANOVA (sex x genotype) and Sidak’s multiple comparisons in B and C. Symbols denote a ***^#^***main effect of genotype in B (***^##^***p < 0.01)) ***^&^***sex(***^&&&^***p<0.0001) in C. Full statistical reporting is available in Table S1. Additional data related to this dataset are available in Figure S4.

### Cnih3 deletion impairs reward-cue association and blunts fentanyl intake during IVSA in both sexes and opioid-seeking during extinction and reinstatement in males only

To assess the impact of Cnih3 deletion on opioid self-administration, mice underwent fentanyl IVSA (Fig 7A). Mice were implanted with jugular vein catheters, then underwent 25 2-hour sessions of fentanyl IVSA during which pokes in a light-cued nose port triggered an infusion of fentanyl (1ug/kg/infusion). The unlit port produced no rewards, but responses were logged to assess preference for the reward-producing port. Next, mice underwent 21 2-hour sessions of extinction in which cues and fentanyl were removed. Last, mice underwent one 2-hour session of cue-induced reinstatement, in which cues, but not the fentanyl reward, were reintroduced to assess drug-seeking behavior. On the first day of fentanyl IVSA, Cnih3 deletion reduced exploratory behavior as assessed by the total number of pokes (Fig 7B). Over the 25 days of IVSA, Cnih3 deletion reduced fentanyl intake (Fig 7C) and the number of infusions self-administered (Fig S5A). Comparison of the average number of infusions between the first and last weeks of fentanyl IVSA showed increases between weeks in WT groups but not KOs (Fig 7D). To assess how intake changed over time, we examined the change in infusions from the second to the last day of fentanyl IVSA and found significant increases within all groups but male Cnih3 KOs (Fig 7E). On the last day of fentanyl IVSA, we compared the number of active (A) and inactive (I) pokes per group to assess preference for the active port at the end of training (Fig 7F), which revealed a lack of preference for the active port only in male Cnih3 KOs (Fig 7F), though there is no difference in discrimination across fentanyl IVSA (Fig S5B). We determined the number of sessions to reach acquisition criteria (2 consecutive sessions where at least 5 infusions are obtained, 70% or more of the pokes are made on the active port, and rewards earned vary by less than 30%), and Cnih3 deletion also impaired opioid-cue association independent of sex (Fig 7G). During extinction, Cnih3 KO reduced opioid-seeking indicated by active nose pokes (Fig 7H), and Cnih3 KOs of both sexes showed no preference for the active lever on the first day (Fig 7I). However, we do not see genotype differences across extinction in discrimination (Fig S5C). Comparing the average number of pokes during the first and last weeks of extinction showed that WTs, but not Cnih3 KOs, significantly reduced responding in the absence of reinforcement (Fig 7J; collapsed due to absence of sex differences). All groups, but Cnih3 KO males showed a decrease in nose pokes from the first to last day of extinction (Fig 7K). Similarly, all groups reinstated drug-seeking behavior except for KO males (Fig 7L). Taken together, Cnih3 deletion blunts fentanyl self-administration in sex specific ways. It appears that, though both sexes initially differ in similar ways from their WT conspecifics, female KOs discriminate the active poke on the last day of fentanyl IVSA and poke more during cue-induced reinstatement than the last day of extinction, while male KOs do not. This suggests that after acquiring IVSA, female KOs begin to equal the self-administration behavior of WT mice while male KOs do not.

**Figure 7:**
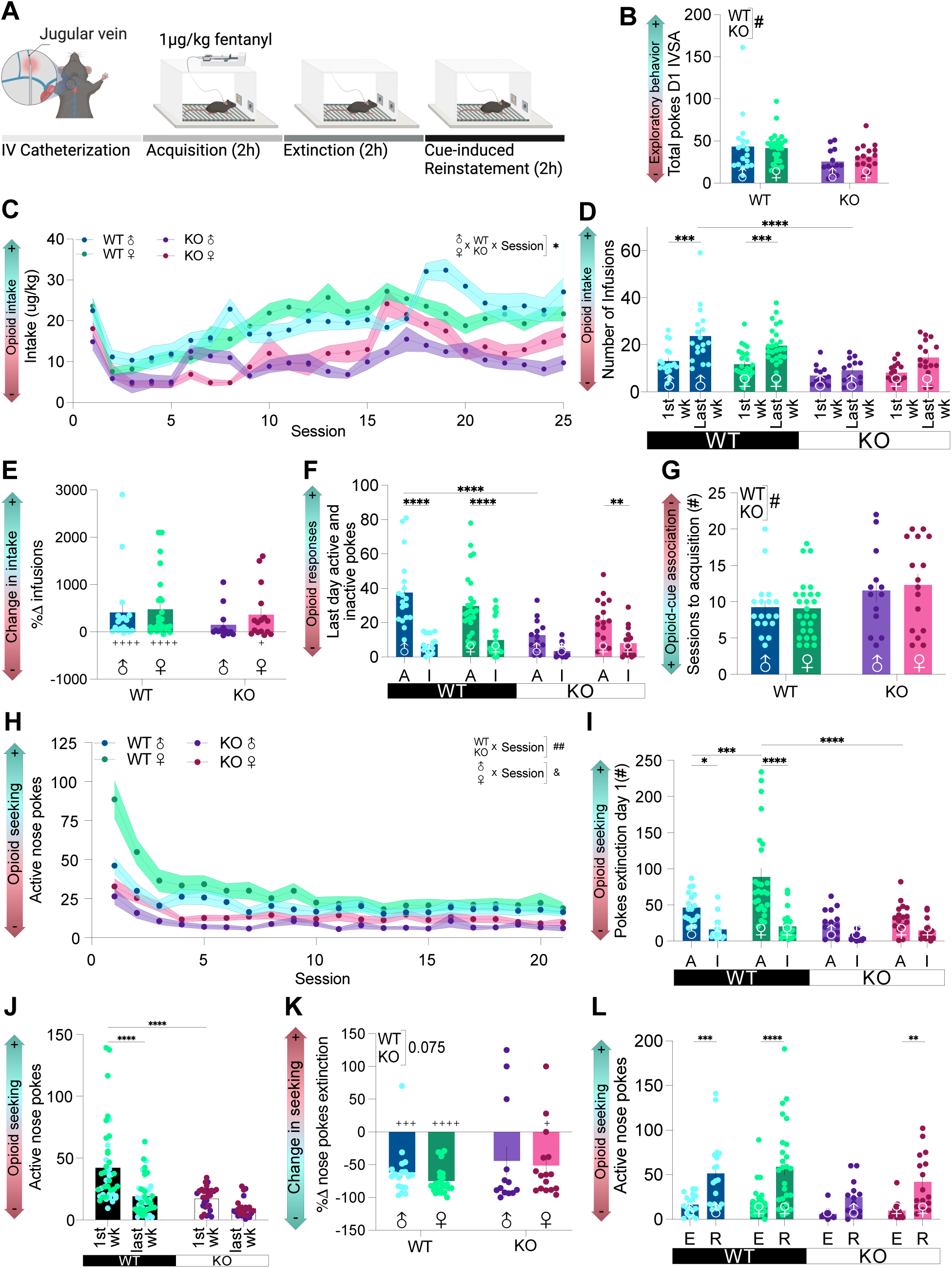
Cnih3 deletion impairs reward-cue association and blunts fentanyl intake during IVSA in both sexes and opioid-seeking during extinction and reinstatement in males only. A. Experimental schematic of fentanyl IVSA experiments. B. Exploratory behavior as assessed by total pokes on the first day of IVSA. C. Opioid intake (ug/kg) during each session. D. Average number of daily fentanyl infusions during the first and last weeks of fentanyl IVSA. E. Percent change in infusion number from day two to the last day of IVSA. F. Active (A) and inactive (I) pokes on the last day of fentanyl IVSA. G. Opioid-cue association as measured by the number of sessions to reach acquisition criteria. H. Opioid seeking as measured by the number of active nose pokes across extinction. I. Opioid seeking as measured by the number of active (A) and inactive (I) pokes on the first day of extinction. J. Average number of active nose pokes during the first and last weeks of extinction. K. Percent change in the number of active pokes from the first to the last day of extinction. L. Opioid-seeking during reinstatement as measured by the number of active nose pokes during the last day of extinction (E) and cue-induced reinstatement (R). Data are reported as mean ± SEM. Two-way ANOVA (sex x genotype) and Sidak’s multiple comparisons in B, E, G, J, and K. Repeated measures (RM) three-way ANOVA (sex x genotype x time) and Sidak’s multiple comparisons in C, D, F, H, I, and L. One-sample t-test for each group to determine the difference from hypothetical 0 in E and K. Symbols denote ^#^main effect of genotype in B and G (^#^p < 0.05) or an interaction between genotype and session in H (^##^ p< 0.01), and an ^&^ interaction between sex and session in H (^&^ p < 0.05). Symbols denote ^+^difference from hypothetical 0 in E and K (^+^p < 0.05, ^+++^p < 0.001, ^++++^p<0.0001), and denote *interactions between sex and genotype in J or sex, genotype, and session in C, D, F, I, and L, (*p<0.05, **p<0.01, ***p<0.001, ****p<0.0001). Full statistical reporting is available in Table S1, Additional data available in Figure S5.

### Cnih3 deletion produces differences in reward-cue association and fentanyl intake during self-administration

To assess the impact of Cnih3 on opioid IVSA and opioid self-administration, we ran a PCA (as in Fig 4) on the behaviors presented in Figure 7 and observed that KO mice cluster tightly together while WT mice have more individual differences, indicating clear genotype differences in this dataset (Fig 8A). In PC1 loadings, Cnih3 deletion produced positive loadings in WT mice and negative loadings in KO mice (Fig 8B). In PC2, Cnih3 deletion did not significantly impact loading (Fig 8C). The largest loadings, and therefore largest contributors, to genotype-based variance in PC1 corresponded to drug-seeking during extinction and post-learning intake (Fig 8D), while PC2 corresponded to reward-cue association and post-learning intake (Fig 8E). Taken together, these data show that there are clear differences between genotypes in opioid-seeking behavior, rooted in drug-seeking during extinction, post-learning intake, and reward-cue association. Generally, KO mice have negative PC1 loadings, exhibiting long acquisition times, low active and inactive lokes on the first day of fentanyl IVSA, as well as low numbers of rewards during weeks 1-5, including the last day, which also showed low inactive pokes. During extinction, KO mice exhibit low active and inactive pokes on the first day, followed by low active pokes during weeks 1, 2, 3, and 4, and low active and inactive pokes during cue-induced reinstatement.

**Figure 8:**
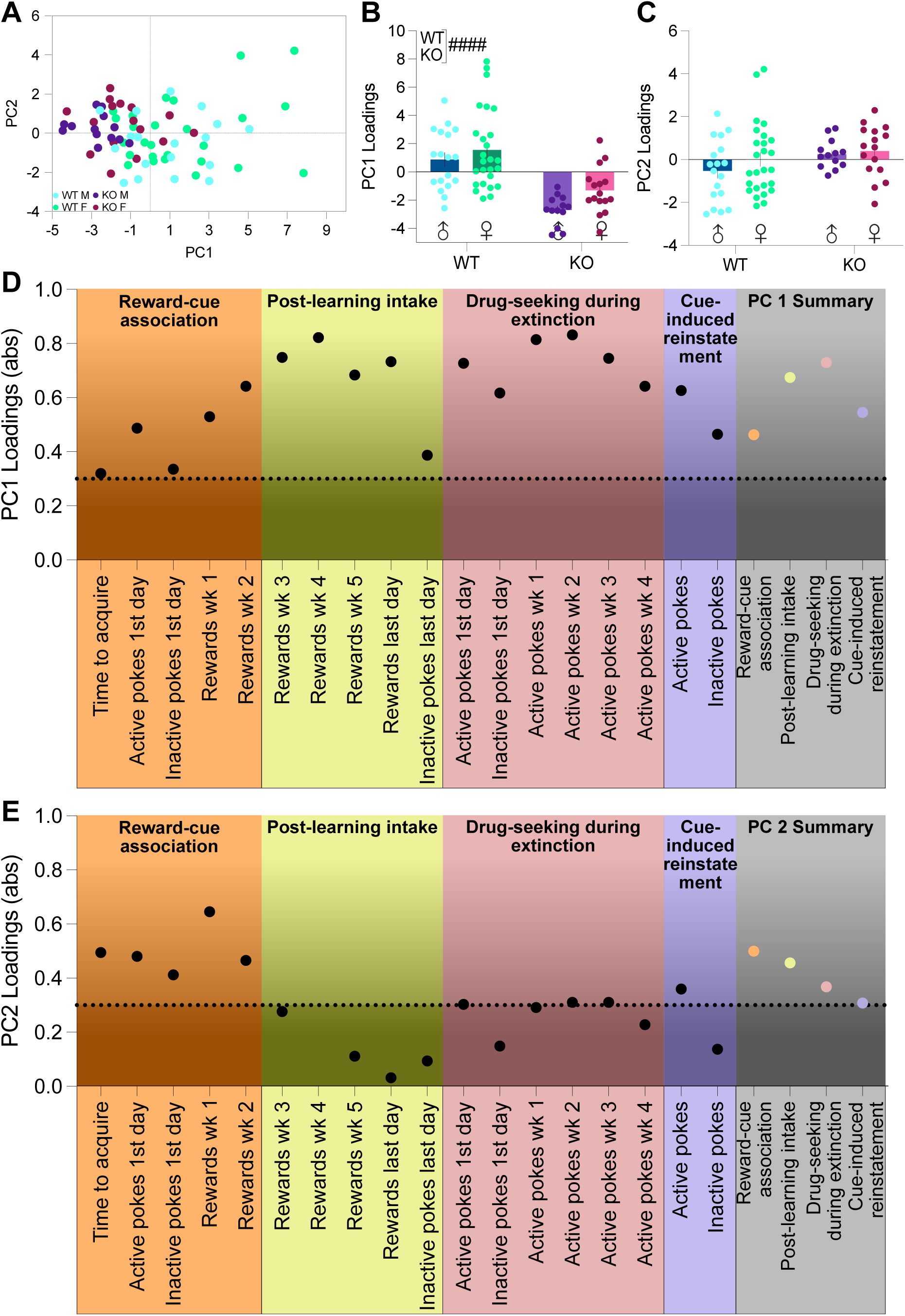
Cnih3 deletion produces differences in reward-cue association and fentanyl intake during self-administration. A. Loadings plot of PC1 and PC2 for each mouse that has completed the experiments described in Figure 8 (n=73). B. PC1 loadings for each mouse. C. PC2 loadings for each mouse. D. PC1 loadings of the variance each described behavior contributes to the dataset. The PC1 summary is the average of the PC loading of all components within a defined type of measure. Loadings above 0.3 are considered significant contributors. E. PC2 loadings of the variance each described behavior contributes to the dataset. The PC2 summary is the average of the PC loading of all components within a defined type of measure. Loadings above 0.3 are considered significant contributors. Data are reported as mean ± SEM in B-C. Average loading of each measure for the dataset shown in D-E. Two-way ANOVA (sex x genotype) in B and C. Symbols denote a ***^#^***main effect of genotype in B (***^####^***p < 0.0001). Full statistical reporting is available in Table S1. Additional data related to this dataset are available in Figure S6.

## Discussion

In this study, we investigated the involvement of CNIH3, an AMPAR auxiliary protein, in opioid use risk factor analysis, sucrose self-administration and reversal learning task, and fentanyl intravenous self-administration. In humans with opioid experience, we found that SNPs in CNIH3 were protective against the progression of OUD, and more protective in women than men. We then employ a mouse model of Cnih3 deletion in mice to comprehensively assess the impact of Cnih3 on risk factors for opioid use. In mice, Cnih3 deletion impaired spatial memory, but not other risk factors such as depression-like behavior and anxiety-like behavior. We then probed the effect of Cnih3 deletion on natural reward seeking and reversal learning using a sucrose self-administration paradigm and found that Cnih3 deletion produced sex-specific deficits in both reward-cue association and reversal learning. Last, we used a fentanyl IVSA paradigm to assess the role of Cnih3 in opioid-seeking and demonstrated that Cnih3 deletion blunted both opioid-cue association and opioid intake. We find that CNIH3 deletion impaired spatial memory, reward-cue association and cognitive flexibility in a sex specific way, and reduced fentanyl self-administration.

### Cnih3 deletion impairs spatial memory, but does not impact other risk factors of OUD

OUD is a complex disorder and several risk factors that predispose people to opioid misuse-many of which involve alterations in glutamatergic transmission, such as pain(*26*), and affective disorders such as anxiety(*18*) and depression(*22*). We assessed the role of Cnih3 in behaviors that constitute risk factors for OUD. Despite the critical role of glutamatergic plasticity in these processes, Cnih3 deletion did not impact well-being, motor coordination, thermal nociception, social memory, anxiety-like or depression-like behavior. The glutamatergic functions supporting these risk factors instead likely rely on or are compensated by other AMPAR support proteins, such as TARPs, in the absence of Cnih3. Nevertheless, Cnih3 deletion moderately impaired spatial memory in the novel object recognition task, consistent with our previous findings(*38*). Spatial memory is a key part of the formation of the contextual and drug-cue associations that make up opioid-associated memories(*10*). Therefore, SNPs in CNIH3 may protect against OUD by disrupting the formation, maintenance, and/or expression of opioid-associated memory. Interestingly, CNIH3 emerged as nominally associated with executive functioning in a large GWAS, suggesting broad correspondence in humans(*42*).

### Cnih3 deletion-based deficits are compounded in opioid use

Cognition and associative learning are key components of maladaptive opioid use, with craving and relapse often triggered by exposure to drug-associated cues and contexts(*43*). Given that Cnih3 deletion impaired spatial memory, we suspected that Cnih3 was also important for the development of reward-cue associations. To test this, we probed the role of Cnih3 in reward-cue association for a natural reward, sucrose, as well as in spatial reversal, a component of cognitive flexibility. This was supported by our finding that female CNIH3 KO mice take longer to acquire sucrose self-administration. Although female KOs appeared to exhibit impaired discrimination during cue association, both sexes have impairments in some aspects of reversal learning, with males taking longer to reach criteria after repeated rule-change, and females administering seemingly fewer rewards during cue-dissociation. Though our experiments do not assess all aspects of cognition, cognitive flexibility, or spatial memory, these results suggest that Cnih3 moderately impairs reward-cue association and cognitive flexibility.

Cognition is an important component in the development of opioid-cue associations, which greatly influences the onset of maladaptive opioid use. Our studies confirm that Cnih3 is a key player in the formation of opioid-cue association and opioid intake, as deletion delays acquisition and decreases fentanyl intake in our IVSA model. Impairments in cognition that worsen spatial/contextual memory formation affect but do not *prohibit* opioid self-administration. PCA shows that reward-cue association and post-learning fentanyl intake were major contributors to the genotype-based variance in drug-seeking. The deficits in spatial memory and natural reward-cue association in our sucrose self-administration experiments predict the delays in opioid-cue association seen in our IVSA model, but the contribution of post-learning intake and drug-seeking during extinction cannot be overstated. These experiments do not assess potential differences in the rewarding properties of fentanyl or differences in motivation for fentanyl, which may contribute to both associative and post-learning genotype differences. Opioid exposure alone alters AMPARs to favor and stabilize GluA1-containing composition(*28,31*), and morphine activation of dopamine neurons in the VTA is dependent on glutamatergic tone(*44*), suggesting that AMPAR trafficking beyond initial learning is also important for opioid-seeking. We therefore suspect that impaired or under-expressed CNIH3 interrupts AMPAR insertion during the early and late phases of opioid use, leading to both our observed decrease in fentanyl self-administration and the protective effects of CNIH3 SNPs in humans.

Our studies indicate that Cnih3 deletion produces moderate deficits in cognition and blunts fentanyl self-administration; however, they do not address the question of *where* in the brain Cnih3 activity is crucial for these behaviors. Our largest deletion-based differences are in spatial memory and reward-cue associations, which are processes mediated by the hippocampus and prefrontal cortex(*45–48*). Both brain regions are also critical for opioid-associated memory(*49–52*) and exhibit high CNIH3 expression(*38,53*). Previous research from our group showed that Cnih3 deletion alters hippocampal AMPAR subunit composition, leading to impaired LTP, and viral overexpression of Cnih3 in the dHPC showed *improved* spatial memory(*38*). These data suggest that Cnih3-mediated plasticity that underlies both spatial memory and opioid use is likely most important in the HPC and PFC.

### The regulatory role of Cnih3 may be sex-dependent

Our reanalysis of the Comorbidity and Trauma Study sample that contributed to the original GWAS shows that the SNPs in CNIH3 were more protective against the progression to daily opioid injection use in women than in men. These differences were recapitulated in some instances in mice with Cnih3 deletion, with sex differences that increase in number as the complexity of the behavior increases. The first major sex differences observed are in the spatial reversal learning task, where female mice tend to perform slightly worse. During initial reward-cue association, nonexistent or small female deficits seem to be exacerbated by the deletion of Cnih3. During the first reversal (cue disassociation), female mice, regardless of genotype, take longer to reach criteria, and female KO mice trend for taking fewer rewards during the first three days. On the second reversal (cue reassociation) male KO mice are the slowest to acquire, while female knockouts discriminate the worst. These results are consistent with reports that females generally have worse cognitive flexibility(*54*) and lower hippocampal AMPAR insertion than males(*55*).

Interesting sex differences arise during fentanyl IVSA, with male KOs seemingly performing worse than female KOs-the opposite effect we see in the human data. Female KOs have a nonzero percent change in the number of infusions earned across the experiment and favor the active nose port on the last session, while male KOs do not. Additionally, male KOs exhibit blunted cue-induced reinstatement. Given that we do not observe sex differences in fentanyl IVSA in our wild-type mice, but do see impairments predominantly in male KOs, we suspect that Cnih3 function may be sex-dependent.

We do not suspect that the sex differences we see in our knockouts are due to differential levels of Cnih3 expression, as previous research has shown no difference in Cnih3 between male and female mice(*38*). We suspect that CNIH3 activity may differ across estrous cycle phase, given that Cnih3 is highly expressed in brain regions with abundant estrogen receptors(*53*). Previous work has shown that Cnih3-related spatial memory deficits in female mice are estrous-cycle dependent(*38*), and that Cnih3 KO mice show larger differences in hippocampal gene expression across the estrous cycle compared to wild-type mice and male mice(*56*). Given the variability in some of our female Cnih3 KO data and the previous works cited above, we suspect that Cnih3 function is modulated by estrogen.

### Cnih3 deletion significantly blunts opioid use and may be a potential therapeutic target

Our data suggest CNIH3 may be a potential therapeutic target to alleviate opioid use. Integrating PCA into our behavioral studies enabled us to unbiasedly recognize large-scale patterns in our data. As the complexity of the paradigm and the saliency of the reward increased (e.g., spatial memory vs. fentanyl self-administration), Cnih3-deletion-based differences became more evident. This is exemplified by the visual stratification of our PC plots, the increasing proportion of variance provided by the first two PCs, and the larger PC loadings (abs.) affiliated with the included variables in each model. Given that intact AMPAR dynamics are necessary for the main contributors to genotype-based variance in our datasets (memory, reward-cue association, cognitive flexibility, and post-learning fentanyl use), we suspect that CNIH3 is a dominant factor in dictating relevant AMPAR dynamics, particularly in those facilitating high-reward states such as opioid use.

Previous research shows that drugs of abuse (including opioids) alter AMPAR subunit composition(*11,32,34,57,58*), a process that is mediated by CNIH3 in vitro(*14*). Our current findings suggest that SNPs in CNIH3 are protective against the development of daily opioid use in humans, and that global deletion does not significantly impact well-being or risk factors for opioid use, such as anxiety-like and depression-like behavior but does significantly blunt opioid self-administration in mice. We hypothesize that in instances of impaired CNIH3 function (such as a loss-of-function mutation, low gene expression, or absence), the AMPAR dynamics necessary for drug-induced plasticity that facilitates maladaptive drug use are also impaired, *protecting* individuals from the development of OUD. This emphasizes CNIH3 as a target that has the potential to alter AMPAR dynamics that facilitate OUD while maintaining necessary AMPAR function. However, the mechanisms by which CNIH3 produces our observed results, as well as the suggestion that CNIH3 functioning may differ by sex, demand further research before contemplating viability as a treatment.

## Conclusion

OUD continues to strain the US as research struggles to keep up with high relapse and overdose rates. Addressing OUD is difficult due to its nature: predisposition does not always translate to the severity of the disorder, and life experiences, access to medical care, social support, and coexisting conditions can greatly alter OUD expression and development. Genetic tools do not hand us the golden key to unlocking OUD, but how we wield them can greatly sharpen the direction of preclinical research to maximize patient care. Unbiased genetic tools, like the GWAS, arm us with a powerful way to identify biomarkers and candidate genes for the prevention and targeted treatment of OUD. Importantly, global CNIH3 deletion does not impact baseline functioning, affective states, or wholly prohibit learning, memory, or cognitive flexibility. Instead, CNIH3 deletion impairs the progression of opioid use, suggesting its importance in the transition to maladaptive opioid use-a role that can hopefully be manipulated to improve the outcomes of those using and misusing opioids.

## Materials and Methods

### Study Design

Sample sizes were determined by power analysis. Data inclusion/exclusion criteria were defined prior to experiment onset and described in each subsection. Outliers were defined prior to the study as values ± 2 standard deviations from the mean. Values that met these criteria were excluded and reported. Treatments/sides for stimulus presentation were counterbalanced and randomized as described in each subsection. Investigators were blinded to mouse genotype during data collection and analysis.

### Human SNP reexamination of sex effects

To conduct a post-hoc reexamination of sex effects using human data from the Comorbidity and Trauma Study (CATS), which was the impetus for the current investigations, we performed logistical regression analyses that included a model with a sex X genotype interaction term and separate additional examinations that were limited to women and men. The sample, methods, and analyses are all otherwise identical to those of the original report(*9*); the phenotype compared opioid dependence individuals with a history of daily IV use (N=1167) to a combined group (N=161) of recreational opioid users and opioid dependent individuals without a history of daily opioid injection. Consistent with our mouse results, we found evidence of stronger protective effects for the minor alleles of CNIH3 SNPs in women (Table 1); interactions reached significance for the three most highly associated SNPs.

### Mice

The experiments were subdivided into *risk factor analysis*, *sucrose self-administration and reversal learning task,* and *fentanyl intravenous self-administration,* each of which included separate cohorts of mice. Details below. All experiments used adult (8–10-week-old) male and female C57BL6J, WT littermate controls, and Cnih3 knockout mice(*38*). Importantly, we assessed potential effects of rearing across all assays by examining both the wild-type littermates of Cnih3 KO mice (no deletion) and C57 wild-type mice but found no significant differences between these groups (F=0.01257-1.799, p=0.1986-0.9117), indicating a lack of influence of maternal care. One exception was in the tail suspension test, for which comparisons were therefore made to littermate controls (Fig 3J-L; Fig S2C-E). Otherwise, wild-type littermates and C57 mice are collapsed and represented as wild-type (WT) throughout.

### Risk factor analysis

Mice were singly housed in a standard 12:12-hr light cycle and tested during the light phase. The behavioral battery included: (i) open field test, (ii) novel object recognition task, (iii) elevated plus maze, (iv) rotarod, (v) nest building, (vi) social interaction test, (vii) hot plate, and (viii) tail suspension test. Mice were habituated to the testing room 1 hour before each behavioral test. Behaviors were recorded and analyzed using AnyMaze software unless otherwise stated. Mice for this experiment performed all eight tasks (described below) with at least one “rest day” between tasks. Behavioral apparatuses were cleaned with Cavicide between mice.

#### (i) and (ii) Open Field Test (OFT) and Novel Object Recognition Task (NORT)

*The OFT was used* to assess exploratory behavior and anxiety-like behavior, and the NORT was used to assess spatial memory. Both took place in a 50x50x50 cm matte grey box for 3 consecutive days. During each day, mice were gently placed in the center of the box and allowed to freely explore for 10 minutes, and the time and distance in each compartment were recorded. NORT Day 1 was a habituation phase (also termed the open field test; OFT), during which mice freely explored the empty chamber. Day 2 was a familiarization phase, during which two identical objects (A) were placed in two investigation zones 5cm from the walls. In addition to the measurements above obtained with AnyMaze, the time spent interacting with each object was hand-scored. On Day 3, the same object A was placed in one investigation zone of the chamber, while a novel object (B) was placed in the other investigation zone. The positions for objects A and B were alternated so that the location of the novel object was counterbalanced. Objects that produced equal investigation times in C57 mice were chosen to avoid object bias(*59*).

#### (iii) Elevated Plus Maze (EPM)

The EPM was used to assess anxiety-like behavior and consisted of an apparatus with 2 open arms, 2 closed arms (50cm length x 10cm width), and a center compartment (10x10cm). Each mouse was placed gently in the center compartment facing away from the experimenter and was allowed to freely explore the apparatus for 10 minutes. Parameters recorded were time and distance in each compartment.

#### (iv) Rotarod

An accelerating rotarod (Ugo Basile) was used to assess motor coordination and balance(*60*). Briefly, mice received a maximum of 5 training sessions where mice spent 120s walking on the rotarod at a fixed speed of 4 rpm. All mice completed training without falling in five attempts or fewer. One hour after successful completion of the training session, the latency to fall as the rotarod accelerated from 4 to 40 rpm over 5 min was assessed. Five consecutive acceleration trials were performed, with 10 min rest time between each trial.

#### (v) Nest-building

The nest-building task was used to assess naturalistic behavior and well-being and was carried out in the home cage(*61*). Each mouse was given 3g of untorn nestlet in a clean cage. 24 hrs later, the intact portions of the nest were weighed to calculate % of the nestlet that was torn.

#### (vi) Social Interaction Test (SIT)

The social interaction test was used to assess social behavior and was conducted in a 60 x 42 x 22 cm box divided into 3 equal chambers. The center chamber was always empty, and the two extreme chambers contained an inverted wired cup that was magnetically secured to the apparatus floor. For this test, mice were allowed to freely explore the apparatus for 10 minutes on 3 consecutive days. Day 1 was a habituation session in which the mice freely explored the three-compartment chamber with both cups empty. On day 2 (social novelty phase), an unknown age and sex matched C57 mouse was placed under one of the two cups. On day 3 (social preference phase), the mouse from day 2 was placed in the same cup and side of the apparatus as the previous day, while a novel age and sex-matched mouse was placed under the cup on the other side. The side of the novel mouse was counterbalanced between subjects. Time spent in each compartment was assessed with AnyMaze, and time spent investigating each cup was hand-scored.

#### (vii) Hot Plate

Mice were gently placed in the center of a 54°C hot plate apparatus (BIOSEB hot and cold plate, BIO-CHP) and observed for signs of thermal nociception (licking/kicking of paws, jumping, etc.). At the first instance of the above behaviors, or once a maximum time of 45 seconds was reached, mice were removed from the hot plate and placed back in the home cage. Latency (s) to the first response was recorded.

#### (viii) Tail Suspension Test (TST)

The tail suspension test was used to assess depression-like behavior and took place in a matte white PLA 3D printed apparatus (50 cm tall, 23 cm wide). A metal rod was secured to a divot in the top of the box. A 4.5 cm long plastic straw was placed at the base of the tail to prevent climbing, and then mice were suspended by the tail from the metal rod. Time spent mobile and immobile, and the number of bouts of each, were measured over 6 minutes.

### Sucrose self-administration and reversal learning task

Mice were group-housed in a reverse 12:12 hr light cycle and tested during the dark cycle. Mice were placed into MedAssociates chambers and learned, without pre-training, to poke a light-cued nose port (counterbalanced between subjects) to receive one chocolate sucrose pellet (TestDiet chocolate sucrose tab 20MG 1818335(5TUT)) in an FR1 schedule of reinforcement during daily 1-hr sessions. Mice could receive a maximum of 30 sucrose pellets, after which the program terminated. Responses on the active and inactive nose ports were recorded (60-sec timeout). Upon reaching acquisition criteria (30 pellets consumed and ≥70% responses on the active nose poke for 3 consecutive sessions), the active and inactive nose ports were switched such that the unlit nose port was reinforced. The same acquisition criteria were used as a measure of cognitive flexibility under these conditions. After re-reaching acquisition criteria, the operant paradigm was changed yet again to match the original conditions such that the lit nose poke was reinforced with the sucrose pellet reward. Time to reach acquisition on this set of parameters was quantified and used as a measure of cognitive flexibility.

### Fentanyl Intravenous Self-Administration (IVSA)

#### IV surgery

Mice were acclimated to a reverse light cycle for at least one week before surgery. Catheters were prepared ahead of time from a 7 cm long piece of polyurethane tubing (Instech, BTPU-027), and a silicone bulb was placed at 0.9 cm from one end and allowed to dry for at least one hour. Before use, the catheter was dipped in Chlorohexidine to sterilize it, and attached to a catheter base (Instech, VAM1B/25) on the side furthest from the silicone bulb. Mice were anesthetized deeply under isoflurane at 2.5-4% and the skin between the shoulder blades and above the right jugular vein was shaved and scrubbed for surgery. A 1 cm incision was made between the shoulders and above the jugular vein, through which the catheter was passed subcutaneously. The jugular vein was isolated by gross dissection, and a small incision was made in the vein using a 26-G needle. Vein hemostats were used to open the incision and slip the catheter (trimmed if needed) into the jugular vein. Once inside the vein, placement was verified by pulling back on the syringe plunger gently and looking for the presence of blood in the tubing. Then, one suture was tied below, and another above, the silicone bulb to secure the catheter. The neck incision was closed, and the mouse was flipped over. The catheter port was placed subcutaneously, leaving the connector exposed, and sutured into place. Then, the back incision was closed, post-operative drugs were administered according to IACUC protocol, and mice were monitored during recovery in a clean cage before returning to the colony.

After surgery, mice were given post-op care (8mg/kg Baytril and 5mg/kg Carprofen once daily) for at least two days and recovered for at least five days before the onset of behavior. Catheters were kept patent and clean using a heparin/gentamicin solution in saline (dose) and were flushed with 0.3ml of the solution daily before entry into the behavioral chambers. At the end of the experiment, catheter patency was verified by flushing at least 0.3mL of 5% Evans Blue through the catheter after decapitation. If dye is visualized outside the jugular vein after flush (leakage), the data were excluded.

#### Self-Administration Task

Mice were singly housed in a reverse 12:12h light cycle and tested during the dark cycle. Fentanyl IVSA took place in operant boxes (Med Associates) equipped with two nose poke ports. The location of the active port was counterbalanced between animals. A cue light within the active port was illuminated to signal drug availability. Nose pokes in the active port resulted in the delivery of fentanyl (1 μg/kg/infusion, i.v.) followed by an 8s ‘timeout’ period during which reinforcement was withheld and the cue light was turned off. Each session lasted 2h. Mice continued daily training for 25 sessions, during which acquisition of the behavior was monitored (criteria: ≥5 infusions, <30% variability in fentanyl intake, and a discrimination index ≥0.7 (DI; active pokes/total pokes) for 2 consecutive days). Next, mice underwent 21 daily 2-h extinction sessions where cue lights were off, and the drug was not available. 24h after the last extinction session, mice underwent a 2-h test of cue-induced reinstatement where non-reinforced nose pokes in response to the reintroduction of the cue light were measured as a proxy for drug-seeking behavior. Mice that did not reach acquisition criteria within 25 sessions, pulled their catheters, failed the catheter patency test following sacrifice, appeared ill, or lost >20% bodyweight were excluded from the analysis (n= 39).

#### Statistical Analysis

The sample, methods, and analyses for reanalysis of human SNP data are identical to those of the original report(*9*). As described, separate analyses were run for men and women in Plink (logistic regressions with age category and three principal components as covariates). Then, genotypic data for the CNIH3 SNPs were extracted, and logistic regressions were run in SAS 9.2 with similar covariates, along with sex and a sex X genotype interaction term.

GraphPad Prism version 10.4.2 was used for all statistics on mouse behavior. All the experiments were replicated at least twice to prevent nonspecific day/condition effects. Behavioral data were assessed for normality using D’Agostino and Pearson tests and Shapiro–Wilk tests, and then analyzed using two-way (sex × genotype) and three-way (sex × genotype × session) ANOVAs. Šídák’s multiple comparisons tests were used to probe significant interactions. Principal component analyses (PCA) were used to perform unbiased pattern recognition to identify factors contributing to variance. In this case, PCA was used to assess the contribution of aspects of each subdivided experiment (*risk factor analysis*, *sucrose self-administration and reversal learning task, and fentanyl intravenous self-administration)* to the variance in the data attributed to genotypic differences. Only mice that completed each task per its subdivided experiment were included in the PCA. In Prism, data were standardized to have a mean of 0 and a standard deviation of 1, and principal components were selected based on parallel analyses, with the percentile level set at 95% and 1000 simulations. Principal components 1 and 2 (PC1, PC2) were selected as they accounted for the highest contribution of variance in each dataset. For interpretation, parameter loadings onto principal components were examined, wherein those >0.3 (absolute value) were considered significant (as previously described(*62*)). All data are expressed as the mean ± SEM, and significance was set at p < 0.05, α = 0.05; see statistical details in table S1.

## Supporting information

Statistics Table

## Acknowledgments

We thank Dr. Louisa Degenhardt and Professor Nick Martin for their work on the initial 2016 GWAS, which inspired this research; Dr. Hannah Frye, Dr. Nicolas Massaly, Dr. Jessica Higginbotham, Dr. Yolanda Campos-Jurado, Dr. Hannah Harder, and Dr. Jessica Cucinello-Ragland for their technical expertise and theoretical contributions to the conceptualization of this project; and Justin Meyer for breeding the colonies used in these experiments and for general managerial support throughout the experiments. Several schematic representations in this manuscript were prepared using https://BioRender.com.

## Author contributions

Conceptualization: TL, JAM

Data curation: TL, AA, EN

Formal analysis: TL, AA, EN

Funding acquisition: JAM

Investigation: TL, AL, TAA, AP, JJD, AA, EN, JAM

Methodology: TL, AL, TAA, AA, EN, JAM

Supervision: TL, AA, EN, JAM

Validation: TL, AL, TAA, AP, JJD, AA, EN, JAM

Visualization: TL, AA, EN

Writing – original draft: TL and JAM

Writing – review & editing: TL, AL, TAA, AP, JJD, AA, EN, JAM

## Data and materials availability

All data reported and additional information required to reanalyze the data reported in this paper are available from the lead contact upon request.

## References

1. Highlights for the 2024 National Survey on Drug Use and Health.

2. Dydyk AM, Jain NK, Gupta M. Opioid Use Disorder: Evaluation and Management. In: StatPearls [Internet]. Treasure Island (FL): StatPearls Publishing; 2025 [cited 2025 Oct 16]. Available from: http://www.ncbi.nlm.nih.gov/books/NBK553166/

3. Edinoff AN, Martinez Garza D, Vining SP, Vasterling ME, Jackson ED, Murnane KS, et al. New Synthetic Opioids: Clinical Considerations and Dangers. Pain Ther. 2023 Apr;12(2):399–421.

4. Deak JD, Johnson EC. Genetics of substance use disorders: a review. Psychol Med. 51(13):2189–200.

5. Kember RL, Vickers-Smith R, Xu H, Toikumo S, Niarchou M, Zhou H, et al. Cross-ancestry meta-analysis of opioid use disorder uncovers novel loci with predominant effects in brain regions associated with addiction. Nat Neurosci. 2022 Oct;25(10):1279–87.

6. Deak JD, Zhou H, Galimberti M, Levey DF, Wendt FR, Sanchez-Roige S, et al. Genome-wide association study in individuals of European and African ancestry and multi-trait analysis of opioid use disorder identifies 19 independent genome-wide significant risk loci. Mol Psychiatry. 2022 Oct;27(10):3970–9.

7. Zhou H, Rentsch CT, Cheng Z, Kember RL, Nunez YZ, Sherva RM, et al. Association of OPRM1 Functional Coding Variant With Opioid Use Disorder: A Genome-Wide Association Study. JAMA Psychiatry. 2020 Oct 1;77(10):1072–80.

8. Polimanti R, Walters RK, Johnson EC, McClintick JN, Adkins AE, Adkins DE, et al. Leveraging genome-wide data to investigate differences between opioid use vs. opioid dependence in 41,176 individuals from the Psychiatric Genomics Consortium. Mol Psychiatry. 2020 Aug;25(8):1673–87.

9. Nelson EC, Agrawal A, Heath AC, Bogdan R, Sherva R, Zhang B, et al. Evidence of CNIH3 involvement in opioid dependence. Mol Psychiatry. 2016 May;21(5):608–14.

10. Heinsbroek JA, De Vries TJ, Peters J. Glutamatergic Systems and Memory Mechanisms Underlying Opioid Addiction. Cold Spring Harb Perspect Med. 2021 Mar;11(3):a039602.

11. Carlezon WA, Nestler Eric J. Elevated levels of GluR1 in the midbrain: a trigger for sensitization to drugs of abuse? Trends Neurosci. 2002 Dec 1;25(12):610–5.

12. Guo Y, Wang HL, Xiang XH, Zhao Y. The role of glutamate and its receptors in mesocorticolimbic dopaminergic regions in opioid addiction. Neurosci Biobehav Rev. 2009 June 1;33(6):864–73.

13. Schwenk J, Harmel N, Zolles G, Bildl W, Kulik A, Heimrich B, et al. Functional proteomics identify cornichon proteins as auxiliary subunits of AMPA receptors. Science. 2009 Mar 6;323(5919):1313–9.

14. Herring BE, Shi Y, Suh YH, Zheng CY, Blankenship SM, Roche KW, et al. Cornichon proteins determine the subunit composition of synaptic AMPA receptors. Neuron. 2013 Mar 20;77(6):1083–96.

15. Certain N, Gan Q, Bennett J, Hsieh H, Wollmuth LP. Differential regulation of α-amino-3-hydroxy-5-methyl-4-isoxazolepropionic Acid (AMPA) receptor tetramerization by auxiliary subunits. BioRxiv Prepr Serv Biol. 2023 Feb 8;2023.02.07.527516.

16. Cull-Candy SG, Farrant M. Ca2+-permeable AMPA receptors and their auxiliary subunits in synaptic plasticity and disease. J Physiol. 2021 May;599(10):2655–71.

17. Kallarackal AJ, Kvarta MD, Cammarata E, Jaberi L, Cai X, Bailey AM, et al. Chronic stress induces a selective decrease in AMPA receptor-mediated synaptic excitation at hippocampal temporoammonic-CA1 synapses. J Neurosci Off J Soc Neurosci. 2013 Oct 2;33(40):15669–74.

18. Riaza Bermudo-Soriano C, Perez-Rodriguez MM, Vaquero-Lorenzo C, Baca-Garcia E. New perspectives in glutamate and anxiety. Pharmacol Biochem Behav. 2012 Feb 1;100(4):752–74.

19. Hasler G, van der Veen JW, Tumonis T, Meyers N, Shen J, Drevets WC. Reduced Prefrontal Glutamate/Glutamine and γ-Aminobutyric Acid Levels in Major Depression Determined Using Proton Magnetic Resonance Spectroscopy. Arch Gen Psychiatry. 2007 Feb 1;64(2):193–200.

20. Duric V, Banasr M, Stockmeier CA, Simen AA, Newton SS, Overholser JC, et al. Altered expression of synapse and glutamate related genes in post-mortem hippocampus of depressed subjects. Int J Neuropsychopharmacol. 2013 Feb;16(1):69–82.

21. He JG, Zhou HY, Wang F, Chen JG. Dysfunction of Glutamatergic Synaptic Transmission in Depression: Focus on AMPA Receptor Trafficking. Biol Psychiatry Glob Open Sci. 2022 Mar 8;3(2):187–96.

22. Ma H, Li C, Wang J, Zhang X, Li M, Zhang R, et al. Amygdala-hippocampal innervation modulates stress-induced depressive-like behaviors through AMPA receptors. Proc Natl Acad Sci U S A. 2021 Feb 9;118(6):e2019409118.

23. Menard J, Treit D. Intra-septal infusions of excitatory amino acid receptor antagonists have differential effects in two animal models of anxiety. Behav Pharmacol. 2000 Apr;11(2):99.

24. Kapus GL, Gacsályi I, Vegh M, Kompagne H, Hegedűs E, Leveleki C, et al. Antagonism of AMPA receptors produces anxiolytic-like behavior in rodents: Effects of GYKI 52466 and its novel analogues. Psychopharmacology (Berl). 2008 June 1;198(2):231–41.

25. Vialou V, Robison AJ, LaPlant QC, Covington HE, Dietz DM, Ohnishi YN, et al. ΔFosB in brain reward circuits mediates resilience to stress and antidepressant responses. Nat Neurosci. 2010 June;13(6):745–52.

26. Cabañero D, Baker A, Zhou S, Hargett GL, Irie T, Xia Y, et al. Pain after Discontinuation of Morphine Treatment Is Associated with Synaptic Increase of GluA4-Containing AMPAR in the Dorsal Horn of the Spinal Cord. Neuropsychopharmacology. 2013 July;38(8):1472–84.

27. Chen T, Wang W, Dong YL, Zhang MM, Wang J, Koga K, et al. Postsynaptic insertion of AMPA receptor onto cortical pyramidal neurons in the anterior cingulate cortex after peripheral nerve injury. Mol Brain. 2014 Oct 31;7:76.

28. Morón JA, Gullapalli S, Taylor C, Gupta A, Gomes I, Devi LA. Modulation of Opiate-Related Signaling Molecules in Morphine-Dependent Conditioned Behavior: Conditioned Place Preference to Morphine Induces CREB Phosphorylation. Neuropsychopharmacology. 2010 Mar;35(4):955–66.

29. Harris GC, Wimmer M, Byrne R, Aston-Jones G. Glutamate-associated plasticity in the ventral tegmental area is necessary for conditioning environmental stimuli with morphine. Neuroscience. 2004 Jan 1;129(3):841–7.

30. Morón JA, Abul-Husn NS, Rozenfeld R, Dolios G, Wang R, Devi LA. Morphine Administration Alters the Profile of Hippocampal Postsynaptic Density-associated Proteins: A Proteomics Study Focusing on Endocytic Proteins*. Mol Cell Proteomics. 2007 Jan 1;6(1):29–42.

31. Billa SK, Liu J, Bjorklund NL, Sinha N, Fu Y, Shinnick-Gallagher P, et al. Increased Insertion of Glutamate Receptor 2-Lacking α-Amino-3-hydroxy-5-methyl-4-isoxazole Propionic Acid (AMPA) Receptors at Hippocampal Synapses upon Repeated Morphine Administration. Mol Pharmacol. 2010 May;77(5):874–83.

32. Xia Y, Portugal GS, Fakira AK, Melyan Z, Neve R, Lee HT, et al. Hippocampal GluA1-Containing AMPA Receptors Mediate Context-Dependent Sensitization to Morphine. J Neurosci. 2011 Nov 9;31(45):16279–91.

33. Hearing MC, Jedynak J, Ebner SR, Ingebretson A, Asp AJ, Fischer RA, et al. Reversal of morphine-induced cell-type–specific synaptic plasticity in the nucleus accumbens shell blocks reinstatement. Proc Natl Acad Sci U S A. 2016 Jan 19;113(3):757–62.

34. Van den Oever MC, Goriounova NA, Wan Li K, Van der Schors RC, Binnekade R, Schoffelmeer ANM, et al. Prefrontal cortex AMPA receptor plasticity is crucial for cue-induced relapse to heroin-seeking. Nat Neurosci. 2008 Sept;11(9):1053–8.

35. Russell SE, Puttick DJ, Sawyer AM, Potter DN, Mague S, Carlezon WA, et al. Nucleus Accumbens AMPA Receptors Are Necessary for Morphine-Withdrawal-Induced Negative-Affective States in Rats. J Neurosci. 2016 May 25;36(21):5748–62.

36. Madayag AC, Gomez D, Anderson EM, Ingebretson AE, Thomas MJ, Hearing M. Cell-type and region-specific nucleus accumbens AMPAR plasticity associated with morphine reward, reinstatement, and spontaneous withdrawal. Brain Struct Funct. 2019 Sept;224(7):2311–24.

37. Rasmussen K, Vandergriff J. The selective iGluR1-4 (AMPA) antagonist LY300168 attenuates morphine-withdrawal-induced activation of locus coeruleus neurons and behavioural signs of morphine withdrawal. Neuropharmacology. 2003 Jan;44(1):88–92.

38. Frye HE, Izumi Y, Harris AN, Williams SB, Trousdale CR, Sun MY, et al. Sex Differences in the Role of CNIH3 on Spatial Memory and Synaptic Plasticity. Biol Psychiatry. 2021 Dec 1;90(11):766–80.

39. Taylor JL, Samet JH. Opioid Use Disorder. Ann Intern Med. 2022 Jan 18;175(1):ITC1–16.

40. Nazarian A, Negus SS, Martin TJ. Factors Mediating Pain-Related Risk for Opioid Use Disorder. Neuropharmacology. 2021 Mar 15;186:108476.

41. Webster LR. Risk Factors for Opioid-Use Disorder and Overdose. Anesth Analg. 2017 Nov;125(5):1741–8.

42. Hatoum AS, Morrison CL, Mitchell EC, Lam M, Benca-Bachman CE, Reineberg AE, et al. Genome-wide Association Study Shows That Executive Functioning Is Influenced by GABAergic Processes and Is a Neurocognitive Genetic Correlate of Psychiatric Disorders. Biol Psychiatry. 2023 Jan 1;93(1):59–70.

43. Vafaie N, Kober H. Association of Drug Cues and Craving With Drug Use and Relapse. JAMA Psychiatry. 2022 July;79(7):641–50.

44. Jalabert M, Bourdy R, Courtin J, Veinante P, Manzoni OJ, Barrot M, et al. Neuronal circuits underlying acute morphine action on dopamine neurons. Proc Natl Acad Sci U S A. 2011 Sept 27;108(39):16446–50.

45. Kilonzo K, Strahnen D, Prex V, Gems J, van der Veen B, Kapanaiah SKT, et al. Distinct contributions of GluA1-containing AMPA receptors of different hippocampal subfields to salience processing, memory and impulse control. Transl Psychiatry. 2022 Mar 14;12:102.

46. Fritch HA, MacEvoy SP, Thakral PP, Jeye BM, Ross RS, Slotnick SD. The anterior hippocampus is associated with spatial memory encoding. Brain Res. 2020 Apr 1;1732:146696.

47. Xiong CH, Liu MG, Zhao LX, Chen MW, Tang L, Yan YH, et al. M1 muscarinic receptors facilitate hippocampus-dependent cognitive flexibility via modulating GluA2 subunit of AMPA receptors. Neuropharmacology. 2019 Mar 1;146:242–51.

48. Stefani MR, Moghaddam B. Rule learning and reward contingency are associated with dissociable patterns of dopamine activation in the rat prefrontal cortex, nucleus accumbens, and dorsal striatum. J Neurosci Off J Soc Neurosci. 2006 Aug 23;26(34):8810–8.

49. Assar N, Mahmoudi D, Farhoudian A, Farhadi MH, Fatahi Z, Haghparast A. D1- and D2-like dopamine receptors in the CA1 region of the hippocampus are involved in the acquisition and reinstatement of morphine-induced conditioned place preference. Behav Brain Res. 2016 Oct 1;312:394–404.

50. Wang Y, Zhang H, Cui J, Zhang J, Yin F, Guo H, et al. Opiate-associated contextual memory formation and retrieval are differentially modulated by dopamine D1 and D2 signaling in hippocampal–prefrontal connectivity. Neuropsychopharmacology. 2019 Jan;44(2):334–43.

51. Shen H, Moussawi K, Zhou W, Toda S, Kalivas PW. Heroin relapse requires long-term potentiation-like plasticity mediated by NMDA2b-containing receptors. Proc Natl Acad Sci U S A. 2011 Nov 29;108(48):19407–12.

52. Lasseter HC, Xie X, Ramirez DR, Fuchs RA. Prefrontal Cortical Regulation of Drug Seeking in Animal Models of Drug Relapse. Curr Top Behav Neurosci. 2010;3:101–17.

53. Uhlén M, Fagerberg L, Hallström BM, Lindskog C, Oksvold P, Mardinoglu A, et al. Tissue-based map of the human proteome. Science. 2015 Jan 23;347(6220):1260419.

54. Gargiulo AT, Hu J, Ravaglia IC, Hawks A, Li X, Sweasy K, et al. Sex differences in cognitive flexibility are driven by the estrous cycle and stress-dependent. Front Behav Neurosci. 2022 Aug 4;16:958301.

55. Monfort P, Gomez-Gimenez B, Llansola M, Felipo V. Gender differences in spatial learning, synaptic activity, and long-term potentiation in the hippocampus in rats: molecular mechanisms. ACS Chem Neurosci. 2015 Aug 19;6(8):1420–7.

56. Mulvey B, Frye HE, Lintz T, Fass S, Tycksen E, Nelson EC, et al. <em>Cnih3</em> Deletion Dysregulates Dorsal Hippocampal Transcription across the Estrous Cycle. eneuro. 2023 Mar 1;10(3):ENEURO.0153-22.2023.

57. Cai YQ, Wang W, Hou YY, Zhang Z, Xie J, Pan ZZ. Central Amygdala GluA1 Facilitates Associative Learning of Opioid Reward. J Neurosci. 2013 Jan 23;33(4):1577–88.

58. Xi ZX, Stein EA. Blockade of ionotropic glutamatergic transmission in the ventral tegmental area reduces heroin reinforcement in rat. Psychopharmacology (Berl). 2002 Nov;164(2):144–50.

59. Inayat M, Cruz-Sanchez A, Thorpe HHA, Frie JA, Richards BA, Khokhar JY, et al. Promoting and Optimizing the Use of 3D-Printed Objects in Spontaneous Recognition Memory Tasks in Rodents: A Method for Improving Rigor and Reproducibility. eNeuro. 2021 Sept 29;8(5):ENEURO.0319-21.2021.

60. Montana MC, Cavallone LF, Stubbert KK, Stefanescu AD, Kharasch ED, Gereau RW. The metabotropic glutamate receptor subtype 5 antagonist fenobam is analgesic and has improved in vivo selectivity compared with the prototypical antagonist 2-methyl-6-(phenylethynyl)-pyridine. J Pharmacol Exp Ther. 2009 Sept;330(3):834–43.

61. Deacon RMJ. Assessing nest building in mice. Nat Protoc. 2006;1(3):1117–9.

62. Fee C, Prevot T, Misquitta K, Banasr M, Sibille E. Chronic Stress-induced Behaviors Correlate with Exacerbated Acute Stress-induced Cingulate Cortex and Ventral Hippocampus Activation. Neuroscience. 2020 Aug 1;440:113–29.

