## Supplemental figures for "Cornichon Homolog-3 (Cnih3) deletion impairs spatial memory, reward-cue association, and fentanyl self-administration behavior"

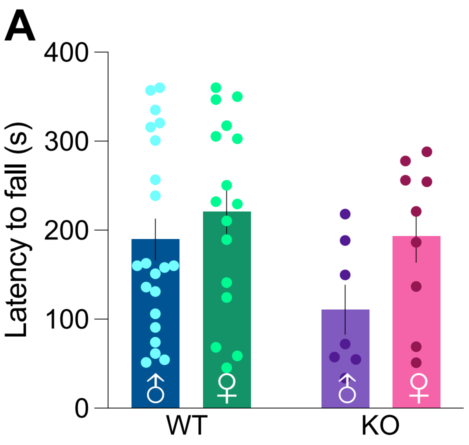


**Figure S1: CNIH3 deletion does not impact motor coordination on Trial 1 (pertains to fig 1)**
A. Motor coordination as measured by latency to fall (s) on the first trial of the accelerating rotarod.

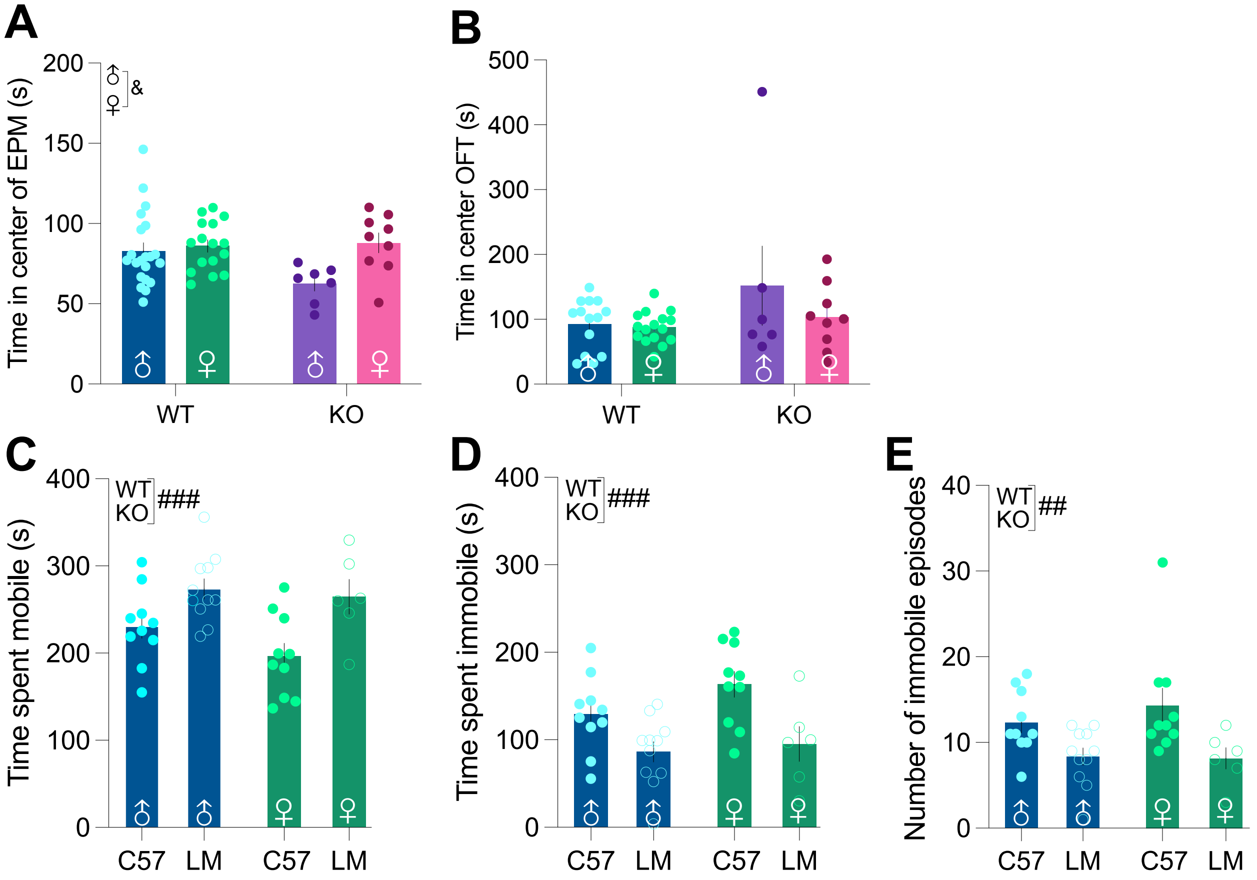


**Figure S2: CNIH3 deletion does not impact time in the center of the EPM or OFT, and WT littermates differ from C57s in depression-like behavior (pertains to fig 3)**

A. Time spent (s) in the center compartment of the elevated plus maze

B. Time spent (s) in the center compartment of the open field test

C. Time spent mobile (s) on the tail suspension test for CNIH3 colony WT littermates and C57 controls

D. Time spent immobile (s) on the tail suspension test for CNIH3 colony WT littermates and C57 controls

E. Number of immobile episodes on the tail suspension test for CNIH3 colony WT littermates and C57 controls

**A**

| Category | Behavior | Variable | PC1 Loading | PC2 Loading |
|---|---|---|---|---|
| Well-being | Nest building | grams of untorn nestlet | 0.088 | **-0.313** |
| | Rotarod | Latency to fall Trial 1 | **0.304** | -0.149 |
| | | Latency to fall Trial 5 | **0.486** | 0.113 |
| | HP | Latency to paw withdrawal | -0.130 | **0.384** |
| Spatial/Social memory | NORT | Time investigating novel object | -0.124 | 0.024 |
| | | Time investigating known object | **-0.382** | -0.204 |
| | | Latency to investigate novel object | **0.728** | -0.075 |
| | SIT | SIT D1 total distance traveled | **0.822** | 0.000 |
| | | SIT D2 time spent with M1 | 0.078 | 0.293 |
| | | SIT D2 time spent with empty | 0.123 | **0.361** |
| | | SIT D3 time spent with M1 | 0.267 | **0.729** |
| | | SIT D3 time spent with M2 | 0.276 | **0.778** |
| Affect | EPM | Distance in open arm | -0.170 | **-0.609** |
| | | Time in open arm | **0.366** | -0.273 |
| | OFT | Time in center zone | 0.292 | **-0.509** |
| | | Distance in center zone | **-0.526** | 0.015 |
| | | Distance in perimeter | **0.760** | -0.190 |
| | TST | Time immobile | 0.210 | -0.019 |
| | | Immobile episodes | 0.254 | **-0.528** |
| | | Latency to first immobile episode | -0.039 | -0.160 |

**B**

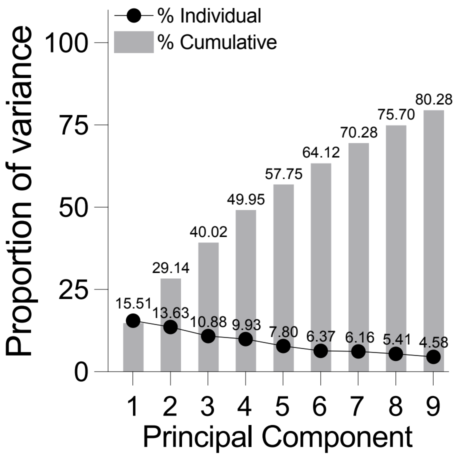


**Figure S3: CNIH3-deletion-based variance in risk factor analysis is minimal (pertains to fig 4)**
A. Variables included in the PCA with loadings for PC 1 and 2. Those less than -0.3 and greater than 0.3 (bold type) are considered significant contributions to the dataset variance
B. Proportion of variance from each principal component. PCs that account for the accumulated 80% of the dataset variance are shown.

**A**

| Category | Experimental Phase | Variable | PC1 Loading | PC2 Loading |
|---|---|---|---|---|
| Reward-cue association | Cue-association phase | Time to acquire | **0.432** | **-0.509** |
| | | Rewards: 1st day | **-0.569** | -0.014 |
| | | Inactive pokes: 1st day | **-0.651** | -0.138 |
| Cognitive flexibility | Cue disassociation and reassociation | Time to acquire: cue disassociation | **0.472** | **-0.351** |
| | | Time to acquire: cue reassociation | **-0.385** | **0.468** |
| | | Rewards: 1st day cue disassociation | **0.460** | -0.132 |
| | | Rewards: 1st day cue reassociation | **0.514** | **0.638** |
| Cognitive inflexibility | Cue disassociation and reassociation | Inactive pokes: 1st day cue disassociation | **-0.613** | **-0.428** |
| | | Inactive pokes: 1st day cue reassociation | 0.117 | **-0.535** |

**B**

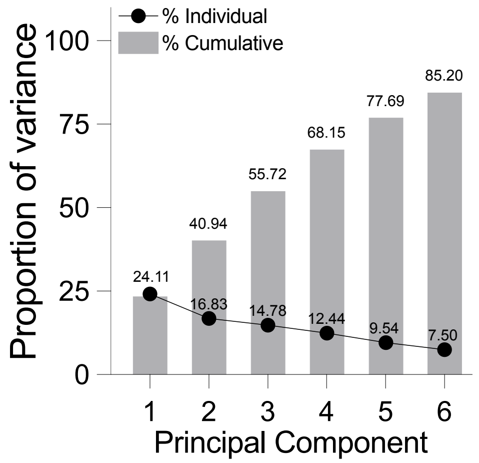


**Figure S4: CNIH3-deletion-based variance in sucrose self-administration and reversal learning is moderate (pertains to fig 6)**

A. Variables included in the PCA with loadings for PC 1 and 2. Those less than -0.3 and greater than 0.3 (bold type) are considered significant contributions to the dataset variance

B. Proportion of variance from each principal component. PCs that account for the accumulated 80% of the dataset variance are shown.

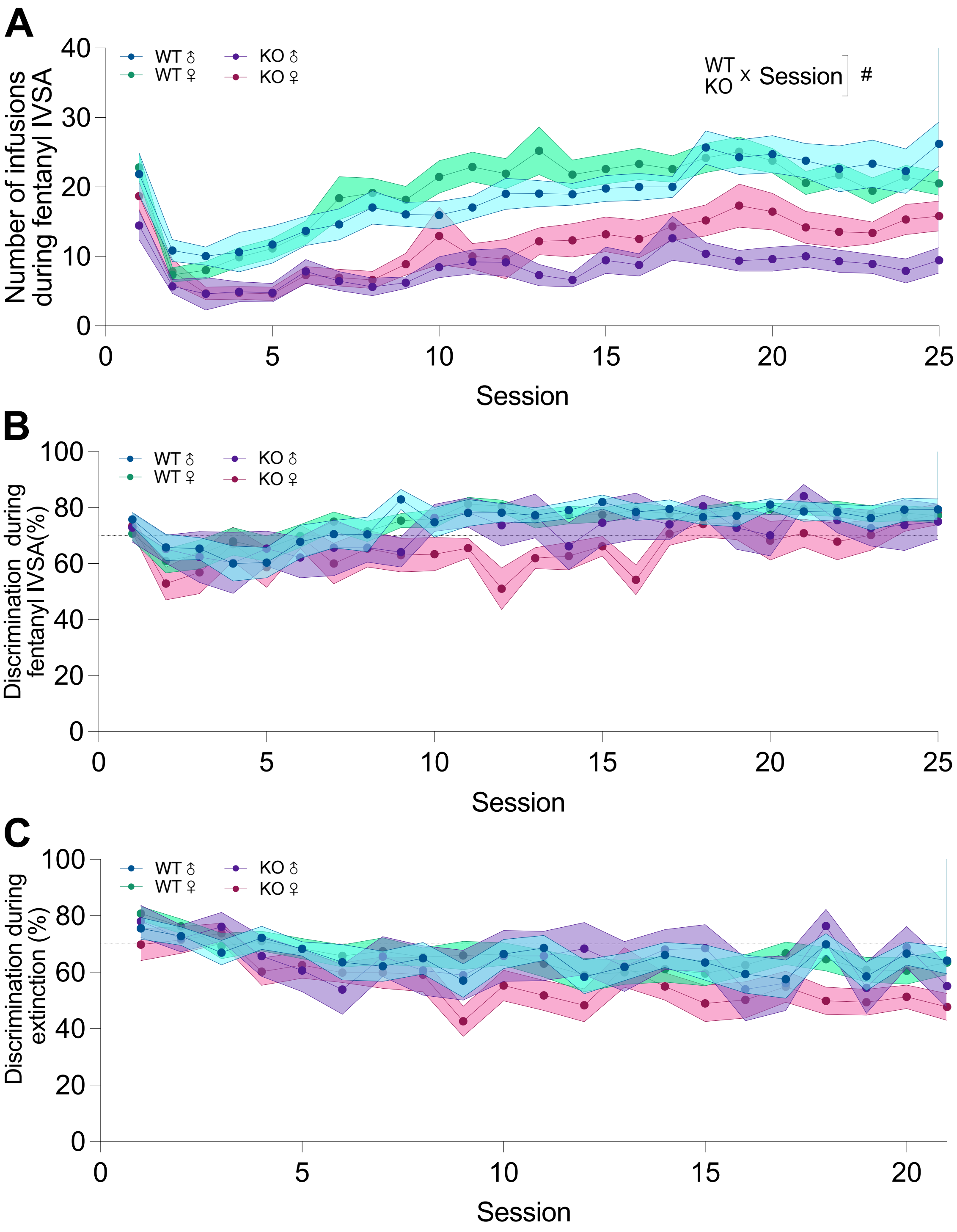


**Figure S5: CNIH3 deletion impairs active nose pokes, but does not impact discrimination for the active nose poke during fentanyl IVSA or extinction (pertains to fig 7)**

A. Average number of infusions earned per session across fentanyl IVSA

B. Average discrimination for the active nose poke (%) across fentanyl IVSA. Discrimination calculated by ((active pokes)/(active pokes + inactive pokes))*100

C. Average discrimination for the active nose poke (%) across extinction. Discrimination calculated by ((active pokes)/(active pokes + inactive pokes))*100

**A**

| Category | Experimental Phase | Variable | PC1 Loading | PC2 Loading |
|---|---|---|---|---|
| Reward-cue association | Acquisition | Time to acquire | **-0.319** | **0.494** |
| | | Active pokes 1st day | **0.487** | **-0.481** |
| | | Inactive pokes 1st day | **0.336** | **-0.412** |
| | | Rewards wk 1 | **0.529** | **-0.645** |
| | | Rewards wk 2 | **0.642** | **-0.465** |
| Post-learning intake | Maintenance | Rewards wk 3 | **0.749** | -0.277 |
| | | Rewards wk 4 | **0.822** | -0.042 |
| | | Rewards wk 5 | **0.683** | -0.112 |
| | | Rewards last day | **0.733** | -0.030 |
| | | Inactive pokes last day | **0.387** | 0.094 |
| Drug-seeking during extinction | Extinction | Active pokes 1st day | **0.727** | **0.303** |
| | | Inactive pokes 1st day | **0.617** | 0.148 |
| | | Active pokes wk 1 | **0.815** | 0.291 |
| | | Active pokes wk 2 | **0.832** | **0.310** |
| | | Active pokes wk 3 | **0.746** | **0.310** |
| | | Active pokes wk 1 | **0.642** | 0.228 |
| Reinstatement | Reinstatement | Active pokes | **0.626** | **0.359** |
| | | Inactive pokes 1st day | **0.464** | 0.137 |

**B**

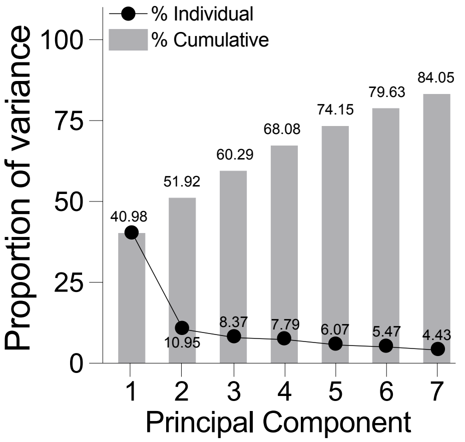


**Figure S6: CNIH3-deletion-based variance in fentanyl intravenous self-administration is large (pertains to fig 8)**

A. Variables included in the PCA with loadings for PC 1 and 2. Those less than -0.3 and greater than 0.3 (bold type) are considered significant contributions to the dataset variance

B. Proportion of variance from each principal component. PCs that account for the accumulated 80% of the dataset variance are shown.
